# A key role for chromosome compartment interactions in directing Xist RNA localisation

**DOI:** 10.64898/2026.05.19.726226

**Authors:** Heather Coker, George Lister, Gabriele Migliorini, Guifeng Wei, Clelia Accalai, Lisa Rodermund, James Davies, Lothar Schermelleh, Neil Brockdorff

## Abstract

X-chromosome inactivation, the mechanism for dosage compensation in mammals, is orchestrated by the long non-coding RNA Xist which localises across the X chromosome in cis. The basis for in cis localisation remains poorly understood. To investigate *in situ* Xist localisation, we established MCPH1-deficient cell models that retain compacted interphase chromosomes, enabling visualisation of individualised chromosome territories. We find that Xist RNA is directed to sites around the periphery of compacted chromosome territories, and moreover that these sites correlate closely with A1-sub-compartments on the X chromosome. We further show that peripheral positioning of A1-sub-compartments occurs on all chromosomes and is a hallmark of early prophase. A key role for compartment interactions in Xist localisation is further supported by analysis of MCPH1-deficient models where Xist is overexpressed (from the X chromosome or an autosomal Xist transgene), and following depletion of HNRNPU, a key factor that anchors Xist RNA to chromosome territories.

## Introduction

X chromosome inactivation (XCI) is a developmentally regulated process in mammals that ensures equal levels of X-linked gene expression in cells of XX female relative to XY male embryos^1^. The XCI process is orchestrated by *Xist*, an X-linked gene which produces a ∼17 kb non-coding RNA required to silence genes along the length of the X chromosome^2–6^. Xist RNA accumulates and spreads in cis along the chromosome from which it is transcribed, recruiting factors that generate a repressive chromatin state and transcriptional silencing chromosome-wide^7,8^. As such, XCI serves as a powerful model for understanding the function of non-coding RNAs and the molecular mechanisms that underpin gene and genome regulation during differentiation and development.

Recent studies have advanced our understanding of the molecular basis for X-linked gene silencing, notably through the identification of two major pathways initiated by the Xist RNA binding proteins (RBPs) SPEN, which mediates histone deacetylation, and HNRNPK which mediates deposition of the histone modifications H2AK119ub1 and H3K27me3 by Polycomb repressive complexes 1 and 2, respectively^9^. Simultaneous perturbation of these pathways fully abrogates Xist-mediated gene silencing^10^. In parallel, there have been advances towards understanding how Xist RNA localises over the X chromosome territory. Depletion of the Xist RNA-binding proteins (RBPs) HNRNPU^11^ and CIZ1^12^ results in the dispersal of Xist RNA throughout the nucleoplasm, suggesting that both proteins function to maintain the exclusive association of Xist with the inactive X chromosome (Xi). CIZ1 function is limited to differentiated cells. Further, RNA Antisense Purification of Xist followed by high-throughput sequencing (RAP-seq) has identified preferred association sites on the Xi that correlate with chromosomal positions with high 3D-contact frequency with the Xist locus^13^, suggesting a role for 3D-chromosome organisation in the spread of Xist RNA. Additionally, there is evidence that regulatory mechanisms maintain appropriate Xist RNA levels to prevent overspill onto nearby chromosomes. More specifically, N6-methyladenosine (m^6^A) modification of Xist RNA modulates Xist RNA turnover via recruitment of the NEXT complex^14^, and recruitment of the H3K9me3 methyltransferase SETDB1 via nascent Xist transcripts functions to autoregulate transcription across the Xist locus^15^. Finally, it has been suggested that specific Xist RBPs facilitate liquid-liquid phase separation and/or Xist condensate formation to promote Xist RNA spreading along the Xi chromosome^16,17^.

Despite these advances, our understanding of Xist localisation to the Xi chromosome remains incomplete. A key challenge in the field is that imaging-based approaches do not allow discrimination between Xist molecules localised to the Xi and those on nearby chromosomes in the context of diffuse chromatin in the interphase nucleus. In this study, we develop cell models that exploit the knockout of the protein MCPH1, which normally binds to the NCAPG2 subunit of condensin II to inhibit its DNA-binding ability in interphase^18^.

Accordingly, in the absence of MCPH1, chromosomes adopt a prophase-like compacted structure during G1 and G2 phases of the cell cycle. Using MCPH1-deficient models, we analyse Xist localisation over the Xi in undifferentiated and differentiated cells, and following perturbation of key pathways that influence Xist localisation. Our findings provide insights into *cis*-limited spreading of Xist RNA and, in parallel, into principles of chromosome organisation during entry into mitosis.

## Results

### MCPH1-deficient XX mESCs undergo X chromosome inactivation

In MCPH1-deficient (ΔMCPH1) mouse embryonic stem cells (mESCs), interphase chromosomes adopt a partially compacted configuration during both G1 and G2 stages of the cell cycle, with a more typical dispersed chromatin morphology apparent during S-phase^18^. Despite this dramatic remodelling of large-scale chromosome organisation, ΔMCPH1 mESCs are fully viable. We set out to test if ΔMCPH1 XX ES cells could be used as a model to analyse Xist RNA localisation during the establishment of XCI. Thus, we used CRISPR/Cas9 facilitated homologous recombination to engineer deletion of the *Mcph1* gene in a previously described interspecific (*Mus domesticus (129S) x Mus castaneus (Cast))* XX mESC line in which Xist expression on one of the two X chromosomes is driven by a doxycycline-inducible promoter (Fig. 1a). The parental line includes a BglG stem-loop/BglG-Halo fusion protein system to enable fluorescent imaging of Xist from the expressed allele^19^. Validated *Mcph1*^-/-^ XX mESC lines were fully viable with a compacted chromosome morphology evident in around 50% of cells, presumably representing G1 and G2/M cells as opposed to S-phase cells (Fig. 1b,c and see also below).

**Figure 1.**
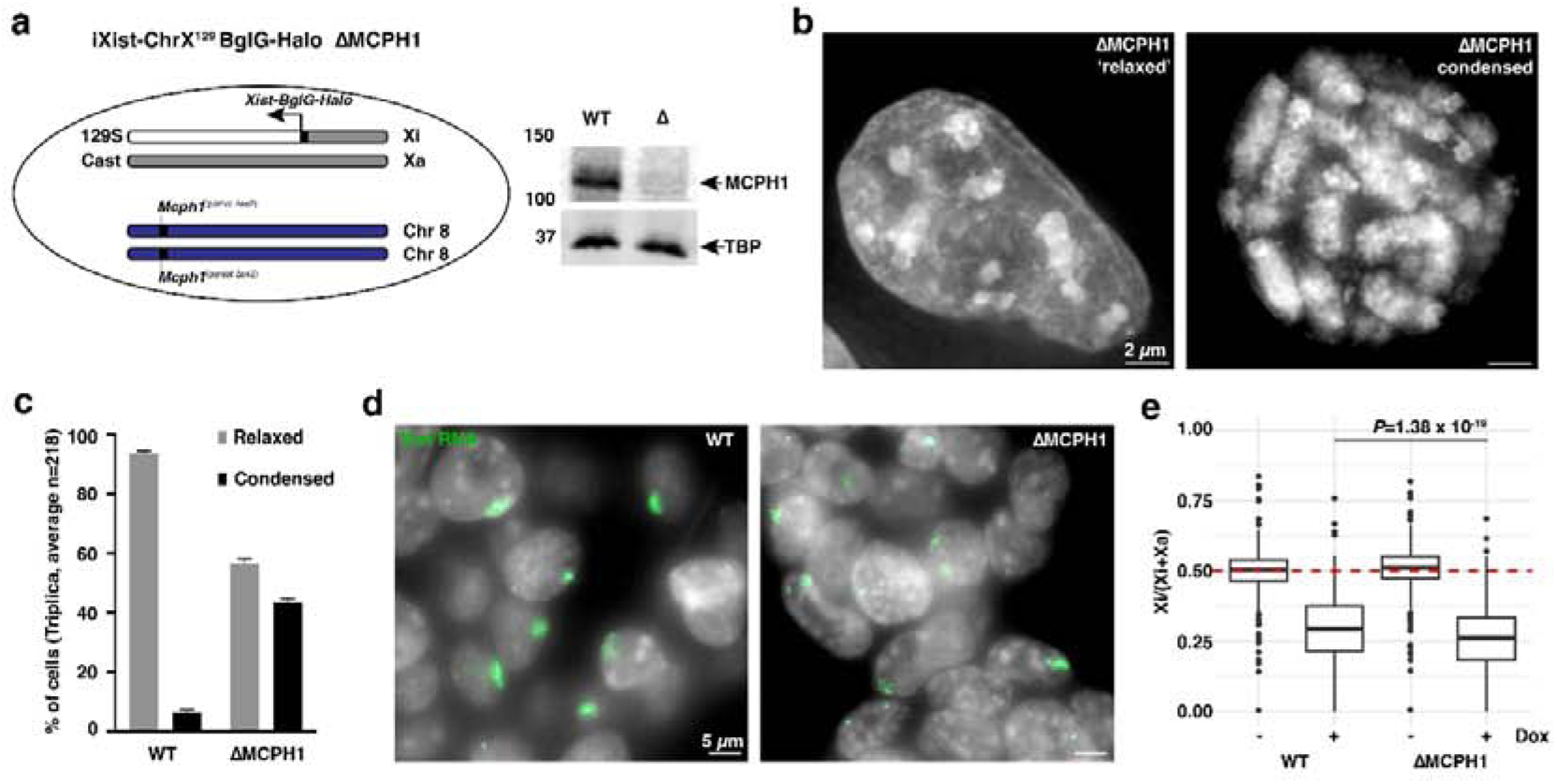
XCI in ΔMCPH1 XX mESCs. **a.** Schematic illustrating the interspecific *Mus domesticus (129S) x Mus castaneus (Cast)* XX mouse ESC line into which a homozygous deletion was engineered into *Mcph1* on chromosome 8 (left). Xist expression was induced from the *129S* X chromosome via rtTA in response to doxycycline (dox) as described^20^. Western blot of WT and ΔMCPH1 cells probed with anti-MCPH1 (and anti-TBP as a loading control) (right). **b.** Relaxed and condensed chromosome morphology in ΔMCPH1 iXist-ChrX^129^ BglG-Halo mESCs cells with DNA counterstained with 4′,6-diamidino-2-phenylindole (DAPI). Maximum intensity projection of deconvolved widefield images. **c**. Scoring of relaxed and condensed chromosome morphology based on DAPI staining in WT and ΔMCPH1 mESCs cells. **d.** Xist RNA FISH (green) in WT and ΔMCPH1 cells showing Xist RNA clouds after induction with doxycycline. DNA counterstained with DAPI (grey), maximum intensity projection image. **e.** ChrRNA-seq^20^ showing silencing of Xi genes is significantly accelerated when Xist expression is induced in the absence of MCPH1 (paired T-test, duplicate experiments, n= 260 genes, WT.

To investigate XCI in ΔMCPH1 XX mESCs, we induced Xist expression with doxycycline for 24 h. RNA FISH analysis revealed discrete Xist RNA domains in the majority of cells (Fig. 1d). Equivalent results were obtained for immunofluorescence (IF) analysis of the repressive chromatin marks H2AK119ub1 and H3K27me3 mediated by Polycomb repressive complexes (PRC) 1 and 2, respectively (Extended Data Fig. 1a,b). Consistent with these observations, X-linked gene silencing as determined by allelic ChrRNA-seq^20^ proceeded normally, albeit at a moderately enhanced rate compared to parental XX mESCs (Fig. 1e). There were no differential effects on genes belonging to distinct sub-classes, for example, based on their relative rate of silencing or position along the X chromosome (Extended Data Fig. 1c,d). Levels of Xist RNA, as determined by analysis of ChrRNA-seq data, were unaffected by MCHP1 deficiency (Extended Data Fig. 1e).

We then used Xist RNA FISH and super-resolution 3D-SIM imaging to examine Xist localisation in ΔMCPH1 mESCs in greater detail^21^. Co-staining for H3S10 phosphorylation enabled identification of cells in the G2/M phase of the cell cycle (Extended Data Fig. 2a-d) with G2 cells exhibiting H3S10P confined to centromeric regions, compared to H3S10P along chromosome arms (whilst retaining an intact nuclear membrane) in early prophase cells. As previously reported^18^, in ΔMCPH1 cells we could further discriminate cells in S-phase (diffuse interphase chromatin) and G1 (compacted chromosomes and absence of H3S10 signal) (Extended Data Fig. 2a). Unlike in wild-type (WT) cells, in which the Xist territory was evident as a uniform ’cloud’ of foci, the ability to visualise individualised chromosome territories in both G1 and G2 ΔMCPH1 cells enabled us to determine that in these cells, most Xist RNA molecules localise to the territory of a single chromosome, the Xi (Fig. 2a-c). Interestingly, Xist localisation on the compacted Xi occurred in a distinctive pattern, with foci restricted to a chromatin layer that circumscribes the chromosome periphery (Fig. 2b,c, Supplementary Movie 1). Individual Xist foci were nevertheless seen to lie in interchromatin spaces rather than entirely outside the discernible Xi chromosome territory (Extended Data Fig.2e), as has been reported previously for WT cells XX cells^21^.

**Figure 2.**
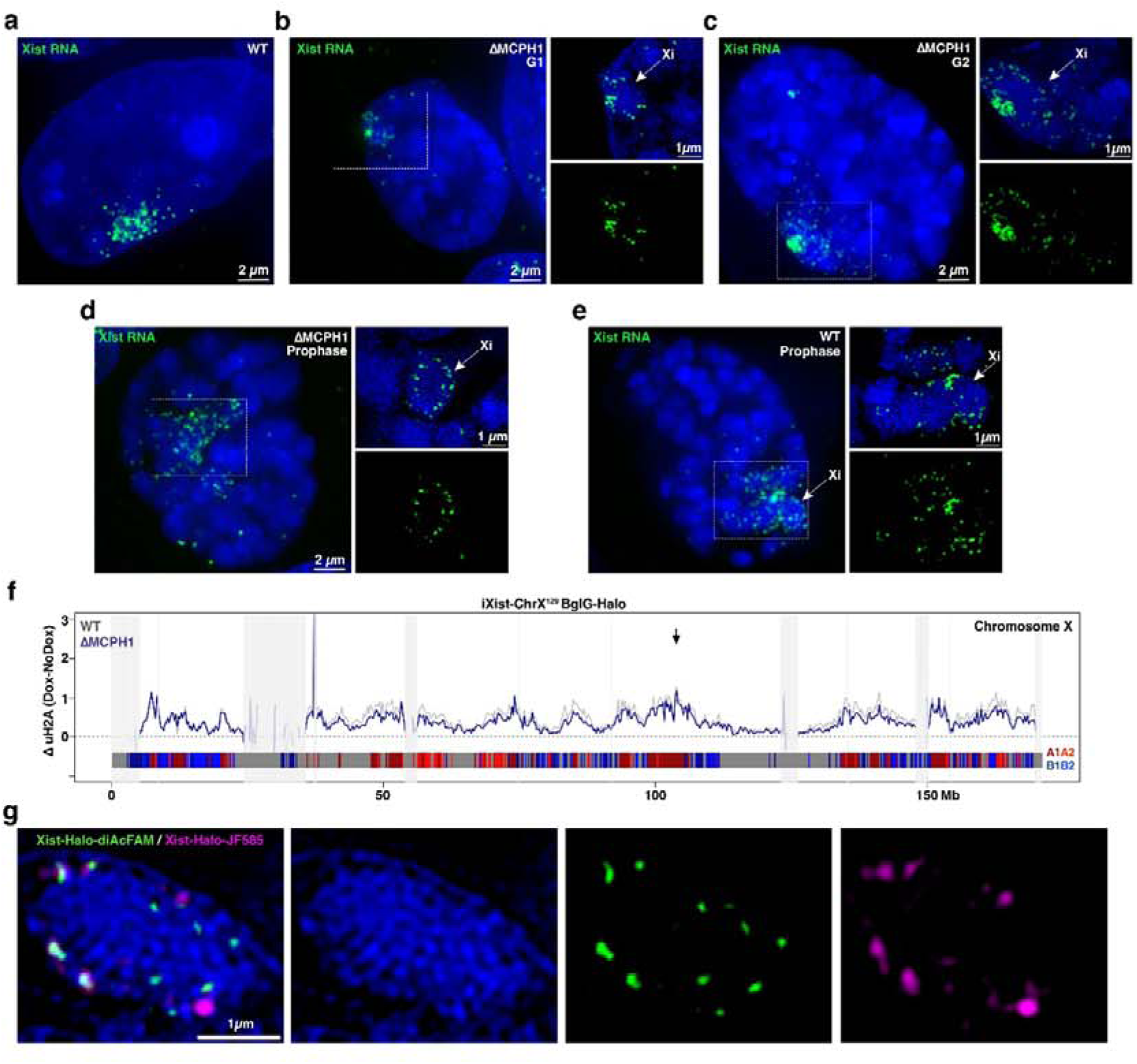
Xist RNA on Xi localises to chromosomal A1-sub-compartments. **a-e**. Xist RNA FISH (green) in iXist-ChrX^129^ BglG-Halo mESC nuclei with DNA counterstained with DAPI (blue). Zooms of Xi regions are shown in insets. **a.** WT cell, **b.** ΔMCPH1 cell in G1 **c.** ΔMCPH1 cell in G2. **d.** ΔMCPH1 cell in prophase. **e**. WT cell in prophase. Cell cycle stage was determined by H3S10P staining (Figure S2). Whole nuclei are maximum intensity projections of deconvolved widefield images and insets show maximum intensity projections of up to 5 3D-SIM z sections. **f.** H2AK119ub1 nChIP profile on the X chromosome from iXist-ChrX^129^ BglG-Halo mESCs showing enrichment of H2AK119ub1 across the X chromosome after 24 h of Xist induction in the parental cell line (grey line, two merged replica experiments, and after MCPH1 deletion (blue line, two merged replica experiments). Blocked out panels (light grey) represent regions/genomic bins where the input signal is significantly different (three standard deviations) from its mean value as computed across the X chromosome (see Materials and Methods). Chromosome sub-compartments depicted in the lower bar: A1 dark red, A2 light red, B1 dark blue, B2 light blue, dark grey; uncertain compartment assignment. The arrow shows the location of the Xist gene. **g.** Representative image of an ΔMCPH1 nucleus with compact chromosomes from RNA-SPLIT experiment demonstrating coupling of temporally separated Xist-Halo diAcFAM (Pulse1) and Xist-Halo JF585 (Pulse 2) molecules on the Xi chromosome. DNA is counterstained with DAPI (blue).

Peripheral Xist localisation was also evident in WT and ΔMCPH1 cells that are in early prophase (Fig. 2d,e). This latter observation indicates that the physiological organisation of Xi during early prophase (i.e. with condensin II but not condensin I loaded) underlies localisation of Xist RNA towards the periphery of the compacted Xi chromosomes.

To further investigate the unusual pattern of Xist localisation over compacted Xi chromosomes we performed allelic H2AK119ub1 native (n) ChIP-seq to define Xist localisation sites^19^. Gain of H2AK119ub1 on Xi (Extended Data Fig. 2f) closely correlates with direct mapping of Xist-bound sites using RAP-seq (Extended Data Fig. 2h )^13,22^. As shown in Fig. 2f, the pattern of Xi specific H2AK119ub1 in ΔMCPH1 mESCs is indistinguishable from that seen in parental XX mESCs, indicating that Xist continues to be bound at the preferred association sites defined in previous studies^13,22^. Interestingly, we observed that preferred association sites align precisely with X chromosome regions defined as A1-sub-compartments in Hi-C analysis^23^ (Fig. 2f, Extended Data Fig. 2g,h). A1-sub-compartments are characterised as having a high gene density and to undergo DNA replication in very early S-phase^23^. Consistent with these features we note correlation of these regions with high SINE repeat density, and anti-correlation with LINE1 (L1) elements (Extended Data Fig. 2g-i). Taken together, these observations imply that A1-sub-compartments are arranged peripherally on compacted Xi chromosomes in ΔMCPH1 cells, and that this mirrors the arrangement of the Xi in normal cells at prophase (see also below).

The BglG stem-loop/BglG-Halo fusion protein system for fluorescent imaging of the expressed Xist allele enabled us to analyse the dynamic behaviour of Xist in ΔMCPH1 cells by using RNA-SPLIT, in which successive waves of Xist RNA are imaged by dual-pulse labelling with Halo-ligands of distinct emission spectra^19^. Our prior analysis using the parental XX mESC line revealed that newly synthesised Xist molecules co-localise with pre-synthesised molecules, a phenomenon that we term Xist RNA coupling. Whilst the underlying basis of Xist coupling is not well understood, we proposed that it relates to a handover mechanism whereby Xist molecules are transferred to and between preferred association sites. RNA-SPLIT performed on the ΔMCPH1 mESCs revealed recapitulation of Xist coupling on compacted Xi chromosomes with co-localised pre-synthesised and newly synthesised molecules, again positioned towards the chromosome periphery (Fig. 2g). This observation indicates that the dynamic behaviour of Xist RNA is broadly unaffected by the compacted chromosome configuration.

### A unique organisation of chromosome compartments in ΔMCPH1 cells

To validate our observations of Xist localisation on Xi in ΔMCPH1 mESCs, we used CRISPR/Cas9-directed mutagenesis to delete the *Mcph1* gene in mouse C127 cells, an XX somatic cell line derived from a mammary gland tumour (Extended Data Fig. 3a). ΔMCPH1 C127 cells were fully viable and had compacted interphase chromosomes (Extended Data Fig. 3b). The proportion of cells with compacted chromosomes was somewhat higher than in ΔMCPH1 mESCs, which likely reflects a smaller proportion of cells in S-phase (Extended Data Fig. 3b,c). RNA-seq analysis revealed a relatively small number of differentially expressed genes (Extended Data Fig. 3d), in accordance with overall cell viability.

The flat morphology of C127 cells facilitates fluorescence imaging of nuclei, allowing improved visualisation of compacted interphase chromosomes. As in mESCs, in WT C127 cells, Xist molecules appear as a ’cloud’ of distinct foci over the Xi territory (Fig. 3a). In ΔMCPH1 cells, analysis of Xi-associated Xist RNA (Fig. 3b, Supplementary Movie 2), or CIZ1 (used herein as a secondary IF marker for Xist RNA), clearly recapitulated the specific localisation to the chromosome periphery (Fig. 3b-e). This pattern was evident in G1, G2 and prophase (Extended Data Fig.3e). A strong enrichment of Polycomb-mediated H3K27me3 was also observed around the Xi periphery (Fig. 3e). We infer from these observations that Xist localisation at preferred association sites (A1-sub-compartments on Xi) is retained during maintenance of XCI in fully differentiated XX somatic cells. Peripheral localisation of Xist, CIZ1 and H3K27me3 was also observed in WT C127 cells at prophase (Fig. 3f, g, Supplementary Movie 3), supporting the proposal that the unique organisation of Xi A1-sub-compartments on compacted chromosomes in ΔMCPH1 cells and during early prophase in normal cells is equivalent.

**Figure 3.**
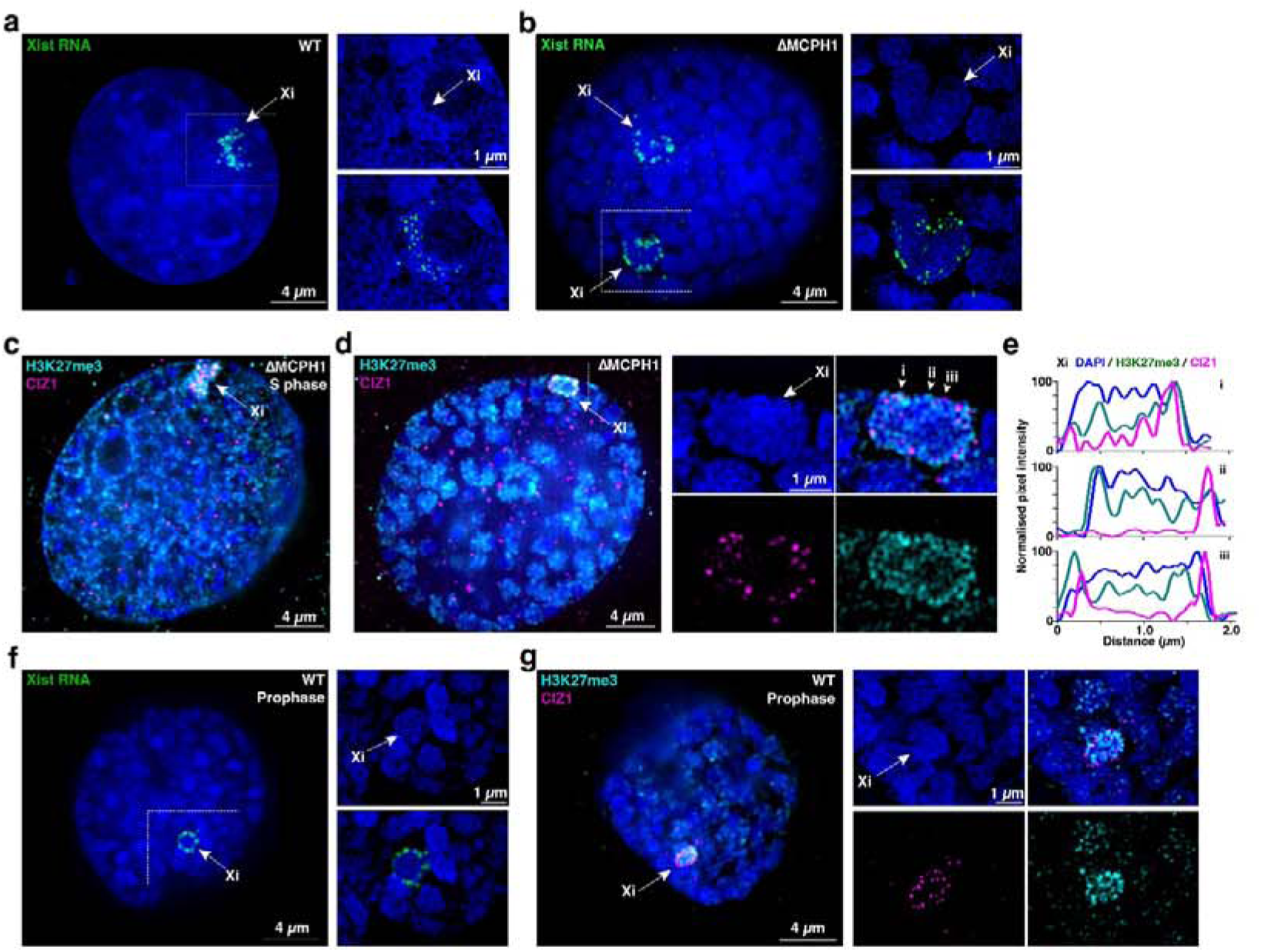
Localisation of Xist RNA in ΔMCPH1 C127 XX somatic cells. **a.** Xist RNA FISH (green) in WT C127 cell and **b.** ΔMCPH1 C127 cell. **c.** IF for H3K27me3 and CIZ1 in C127 ΔMCPH1 cell at S-phase (relaxed chromosome morphology) compared to **d.** C127 ΔMCPH1 cell with condensed chromosome morphology. **e.** H3K27me3 and CIZ1 distributions across the Xi are depicted as IF signal intensity profiles along the chromosome at the lines denoted i-iii in panel **d**, with the start of each line indicated by an arrowhead. **f.** Xist RNA FISH (green) in prophase WT C127 nucleus. **g.** IF for H3K27me3 and CIZ1 in prophase WT C127 nuclei. The WT prophase stage is determined by chromosome morphology. In all panels, white rectangles indicate Xi regions in zoomed views. DNA is counterstained with DAPI (blue). Images of nuclei display maximum intensity projections of widefield deconvolved data, with insets showing maximum intensity projections of 3 3D-SIM z sections.

In normal interphase cells, chromosomes are largely organised so that transcriptionally inactive heterochromatin (B-compartments) preferentially localises towards the nuclear and/or nucleolar periphery and transcriptionally active euchromatin (A-compartments) towards more internal positions (reviewed in^24^). Given that the Xi in ΔMCPH1 cells shows a converse inside-outside configuration, we were interested to determine if this is a general feature of compacted chromosomes in ΔMCPH1 cells and of chromosomes in early prophase. In support of this proposal, IF analysis of markers of actively transcribed euchromatin, specifically elongating RNA Polymerase II (RNA Pol II Ser2P)(Fig. 4a-c) and the histone PTM H3K4me3 (Extended Data Fig.4a), revealed localisation outside or on the periphery of the compacted chromosome territories, except on the transcriptionally silent Xi. H3K27me3, a marker of facultative heterochromatin, on the other hand, localised throughout the compacted chromosome territories (Fig. 4b, Extended Data Fig. 4a), except as noted above on the Xi, where there is additionally strong enrichment around the chromosome periphery. This configuration of chromatin domains was evident in G1, G2 and prophase cells, shown by IF analysis of RNA Pol II Ser2P (Extended Data Fig. 4b) and on early prophase chromosomes in WT C127 cells, shown by IF analysis of H3K4me3 and H3K27me3 (Extended Data Fig. 4c).

**Figure 4.**
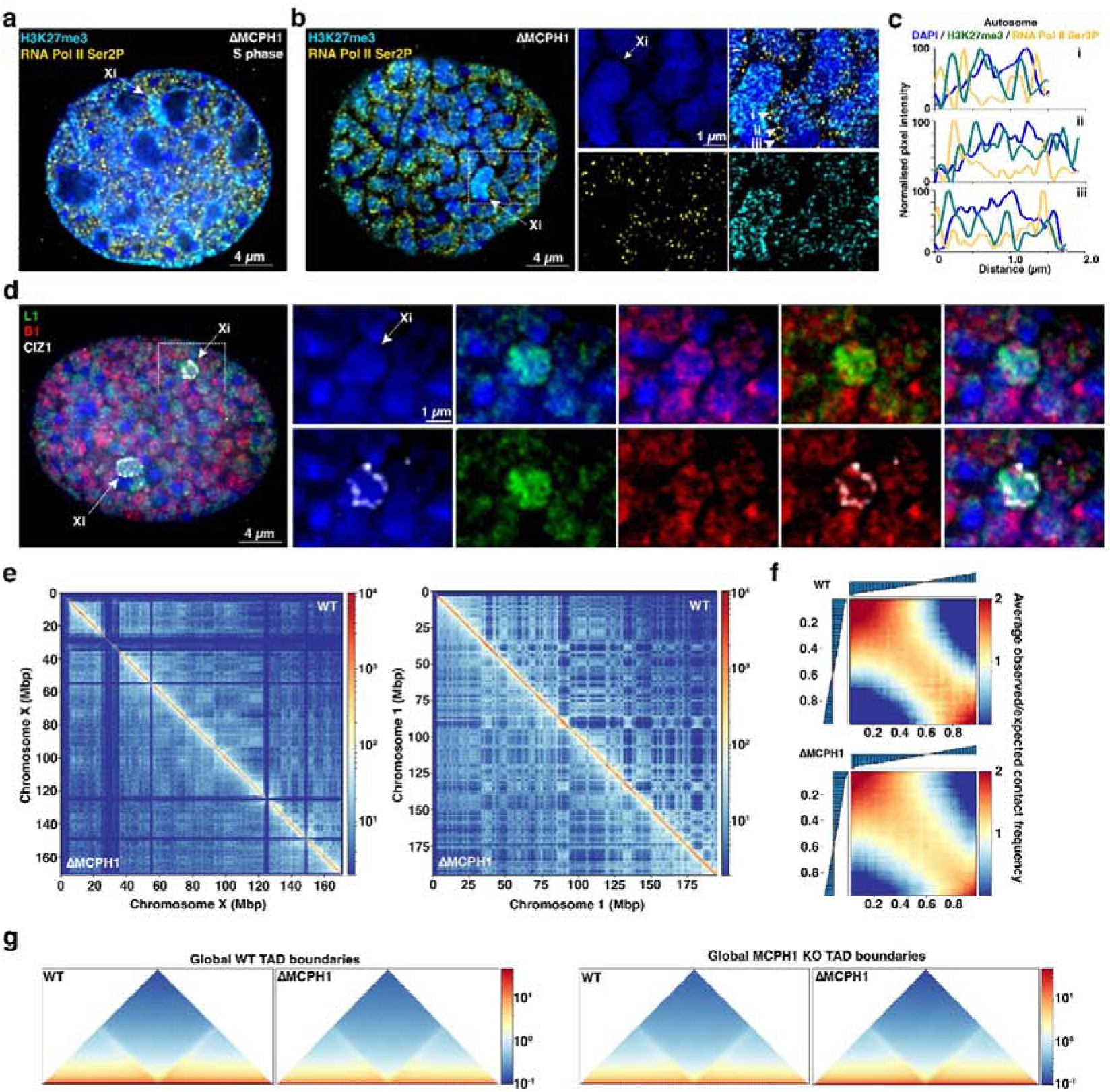
Inside-outside configuration of A- and B-compartments in ΔMCPH1 cells. **a.** IF for H3K27me3 and RNA Pol II Ser2P in a C127 ΔMCPH1 cell at S-phase (relaxed chromosome morphology) compared to **b.** a C127 ΔMCPH1 cell with condensed chromosome morphology. **c.** H3K27me3 and RNA Pol II Ser2P distributions across an autosome are depicted as IF signal intensities along lines i-iii across the chromosome, with the start of each line indicated by an arrowhead. **d.** Immuno-RASER-FISH on a C127 ΔMCPH1 cell with condensed chromosome morphology, with L1 and B1 probes recognising LINE1 and SINE elements respectively, and CIZ1 IF indicating Xist RNA localisation. DNA is counterstained with DAPI (blue). Nuclei are maximum intensity projections of the widefield deconvolved image, with white rectangles indicating the region of the inset zoom view. Insets show maximum intensity projections of 3 3D-SIM z sections (IF), or 3 widefield deconvolved z sections (immuno-RASER-FISH). **e**. Hi-C analysis of C127 WT and ΔMCPH1 cells at 250 Mbp resolution on the X chromosome (left panel) and chromosome 1 (right panel). **f.** Hi-C saddleback plots illustrating the difference in A-A, B-B and A-B compartment interaction strength in WT C127 WT (top panel) and DMCPH1 (lower panel) cells. **g**. Global TAD boundaries in WT and MCPH1 KO C127 cells based on TADs determined from WT cells (left panel) or ΔMCPH1 cells (right panel).

To directly test the spatial configuration of chromosomal A- and B-compartments on compacted chromosomes in MCPH1-deficient cells, we developed a methodology based around RASER-FISH^25^, a minimally disruptive DNA-FISH methodology, to analyse SINE and L1 dispersed repeat families that show highly preferred association with A- versus B-compartments respectively (Extended Data Fig. 4d-g). Analysis of compacted chromosomes in ΔMCPH1 C127 cells (Fig. 4d) revealed that focal signals for SINEs are indeed concentrated around the periphery of all chromosomes, whereas L1 signal localises predominantly through the core of the compacted chromosomes. Co-staining for CIZ1 clearly illustrates co-localisation of Xist RNA with focal SINE signals (corresponding to the A1-sub-compartment) (Fig. 4d).

As an orthogonal approach to investigate chromosome organisation in MCPH1-deficient C127 cells, we performed Hi-C^26^ and subsequently quantified chromosome contacts in the context of topologically associated domains (TADs) and chromosome compartments. This analysis revealed that relative to WT C127 cells, there is little or no change in TAD boundaries and a modest reduction in intra-compartment interactions (A-A, B-B vs A-B) (Fig. 4e-g). A likely explanation for the latter observation is that intra-compartment interactions are somewhat restricted on the relatively rigid compacted chromosomes in ΔMCPH1 cells. These findings are in accordance with prior Hi-C analysis of ΔMCPH1 XY mESCs^18^ and with analysis of 3D chromosome contacts in normal prophase cells^27^.

### Spillover of excess Xist RNA analysed in ΔMCPH1 cells

Recent studies have documented spillover of Xist RNA onto neighbouring chromosomes either as a result of excess levels^28^, or during early stages of X inactivation^29,30^, with the latter being linked to Xist-dependent silencing of specific autosomal loci. We set out to investigate this effect in ΔMCPH1 cells, in which the boundaries of individual chromosome territories can be discerned at interphase. We first investigated XX mESCs engineered to enable depletion of the H3K9me3 methyltransferase SETDB1, previously shown to result in a 4-5-fold increase in Xist RNA levels^15^. Thus, we deleted the *Mcph1* gene in XX mESCs that have doxycycline-inducible Xist expression, and SETDB1 modified with a C-terminal FKBP12^F36V^ degron tag^31^ to enable rapid depletion of the protein on addition of dTAG-13 (Extended Data Fig. 5a). Acute SETDB1 depletion in the iXist-ChrX^Cast^ ΔMCPH1 line resulted in significantly higher levels of Xist RNA and accelerated XCI (Extended Data Fig. 5b), consistent with prior results obtained for the parental line^15^.

We went on to analyse overexpressed Xist RNA by RNA-FISH and IF analysis for CIZ1 (Fig. 5a-f, Extended Data Fig. 5c,d). As expected, Xist RNA in iXist-ChrX^Cast^ ΔMCPH1 S-phase cells prior to SETDB1 depletion appears as a diffuse cloud, similar to WT cells, and in cells with compacted chromosomes we could observe Xist localised at the periphery of the single Xi chromosome (Fig. 5a-c, Extended Data Fig.5c). Following SETDB1 depletion we observed a larger cloud in S-phase cells (Fig. 5d), whilst in cells with condensed chromosomes, Xist RNA localised to the Xi and several adjacent chromosomes (Fig. 5e,f, Extended Data Fig. 5d). Xist molecules on the Xi and adjacent chromosomes localised predominantly to the chromosome peripheries (Fig. 5f). Consistent with these observations, nChIP-seq analysis of Xi specific H2AK119ub1 recapitulates A1-sub-compartment specific enrichment (Fig. 5g). We did not find specific regions on non-Xi chromosomes showing a significant gain of Xist-dependent H2AK119ub1, and moreover analysis of ChrRNA-seq data (Extended Data Fig. 5b) did not identify non-Xi chromosomal regions exhibiting Xist-mediated gene silencing.

**Figure 5.**
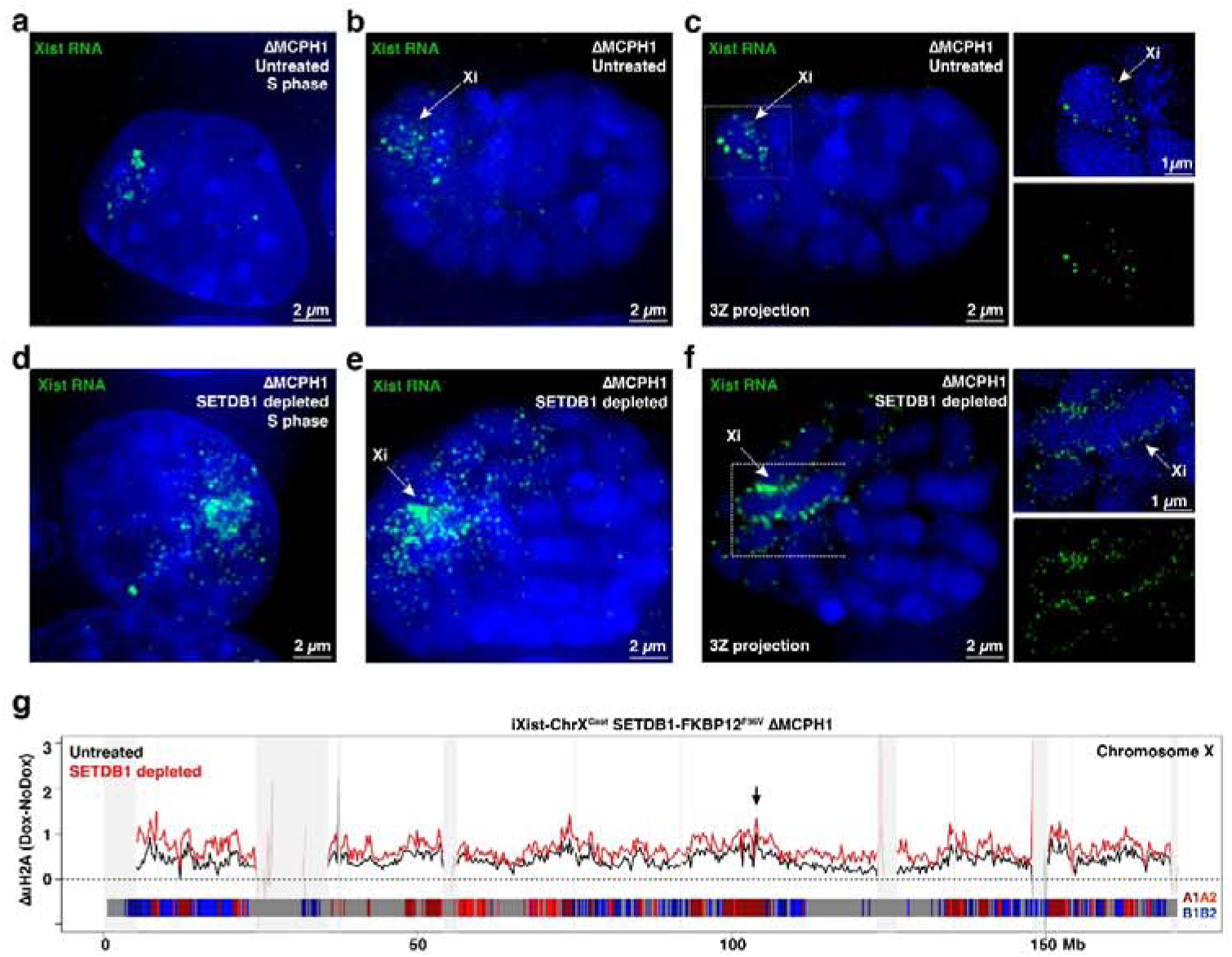
Visualising Xist RNA overspill in ΔMCPH1 cells after SETDB1 depletion. **a, b.** Maximum intensity projection of widefield deconvolved images of Xist RNA FISH (green) in an untreated iXist-ChrX^Cast^ ΔMCPH1 SETDB1-FKBP12^F36V^ cell at S-phase cell (**a**) with relaxed chromosome morphology and an untreated iXist-ChrX^Cast^ ΔMCPH1 SETDB1-FKBP12^F36V^ cell (**b**) with condensed chromosome morphology. **c**. As in **b,** but showing a maximum intensity projection of 3 z-sections of the widefield deconvolved image. Inset zoom (indicated by area in white rectangle) is a 3z projection of the corresponding 3D-SIM image. **d-f**. As in **a-c** for representative cells after depletion of SETDB1 by treatment with dTAG-13. DNA is counterstained with DAPI (blue). **g**. H2AK119ub1 nChIP profile of the X chromosome after 24 h of Xist induction in iXist-ChrX^Cast^ ΔMCPH1 cells in the presence (black line, two merged replica experiments) or absence of SETDB1 (red line, two merged replica experiments). Blocked out panels (light grey) represent regions/genomic bins where the input signal is significantly different (three standard deviations) from its mean value as computed across the X chromosome (see Materials and Methods). Chromosome sub-compartments depicted in lower bar: A1 dark red, A2 light red, B1 dark blue, B2 light blue, dark grey; uncertain compartment assignment. Arrow denotes the location of the Xist gene.

A second model with elevated levels of Xist RNA is iXist-Chr3 XY mESCs^20^, in which Xist expression is driven from a multi-copy doxycycline-inducible transgene located on mouse chromosome 3. In iXist-Chr3 mESCs, Xist-mediated gene silencing is most evident in a region centred on the transgene integration site at the distal end of chromosome 3^20^, and consistent with this, we observe Xist-dependent gain of H2AK119ub1 predominantly in this region (Extended Data Fig. 6a). Interestingly, gain of H2AK119ub1 shows a pattern that is reciprocal to the location of A1-sub-compartments, locating instead to B-compartment regions (Extended Data Fig. 6a). Possibly related to this observation, the Xist transgene on chromosome 3 is integrated into a B-compartment region, whereas the native *Xist* gene on the X chromosome lies within an A-compartment.

To determine patterns of Xist RNA localisation in iXist-Chr3 we generated derivative mESCs with a deletion of *Mcph1* (Extended Data Fig. 6b). Xist RNA FISH and IF analysis for the Polycomb-mediated histone modifications H2AK119ub1 and H3K27me3 and the Xist-associated protein CIZ1 identified large domains following Xist RNA induction (Fig. 6a-c, Extended Data Fig. 6c-e). ChrRNA-seq analysis demonstrated moderately increased silencing of chromosome 3 genes compared to parental iXist-Chr3 cells (Fig. 6f), consistent with observations in ΔMCPH1 XX mESCs (Fig. 1e). Xist RNA levels in iXist-Chr3 cells were largely unaffected by MCPH1 depletion (Extended Data Fig. 6g). In cells with condensed interphase chromosomes, we could observe Xist RNA localising to chromosome 3, and to a significant degree over adjacent chromosomes (Fig. 6b). Close examination indicated that transgenic Xist RNA on chromosome 3 is not strictly confined to the peripheral chromatin layer (Fig. 6b), although Xist molecules associated with adjacent chromosomes were (Fig. 6b).

**Figure 6.**
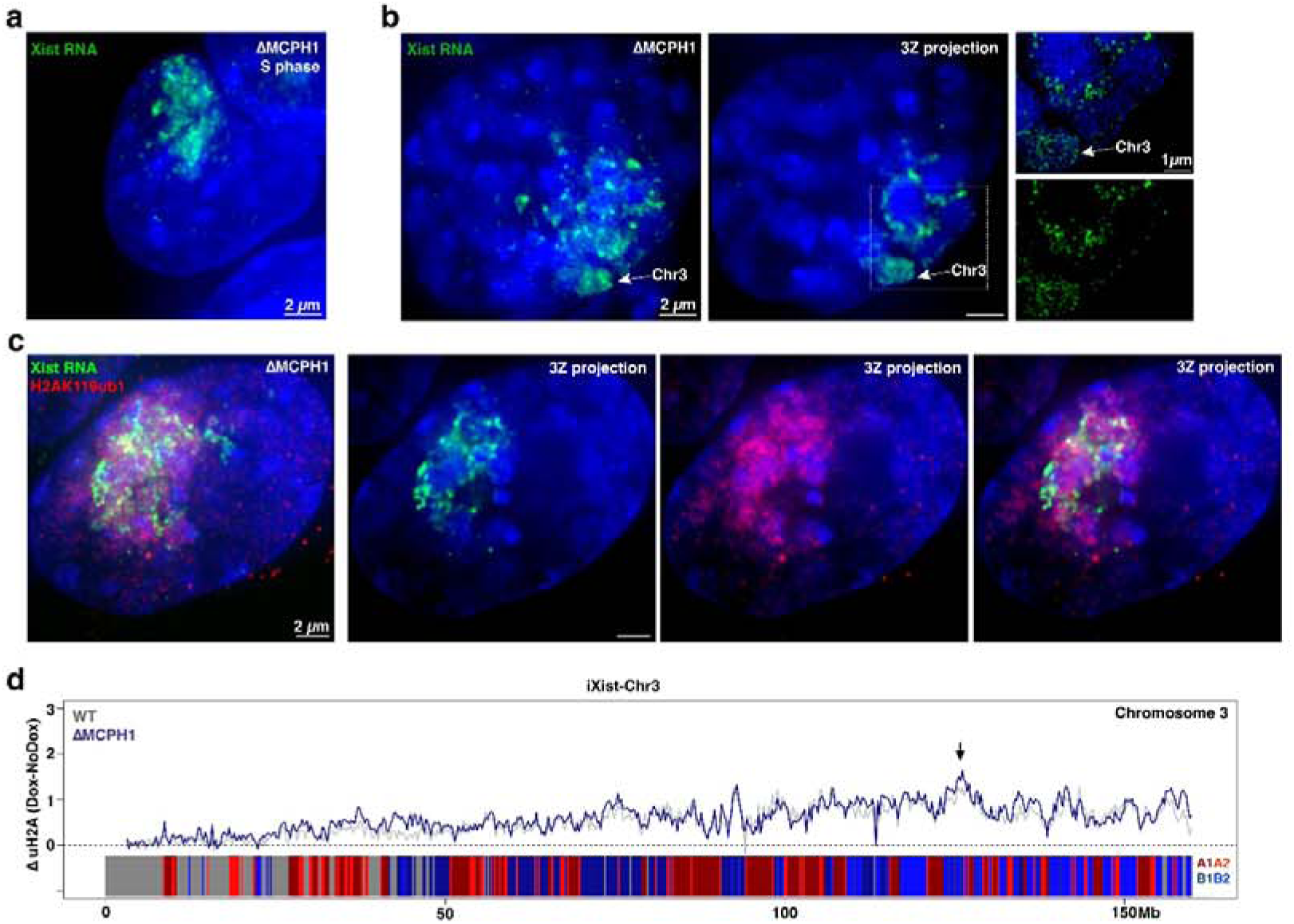
Xist RNA localisation in autosomal transgene iXist-Chr3 ΔMCPH1 cells. Maximum intensity projection of widefield deconvolved images of **a.** Xist RNA FISH (green) in a ΔMCPH1 iXist-Chr3 S-phase cell (relaxed chromosome morphology), and **b**. an ΔMCPH1 iXist-Chr3 cell with condensed chromosome morphology (left panel). The central panel shows a 3z projection of the same widefield deconvolved image. The right panel shows a 3z projection of the area demarcated by a white rectangle, imaged with 3D-SIM. **c**. Example of Xist Immuno-FISH for H2AK119ub1 and Xist RNA in ΔMCPH1 iXist-Chr3 cell with condensed chromosome morphology. The left panel shows a maximum intensity projection of the deconvolved widefield image, with subsequent panels illustrating colour-channel splits of 3z projections of the deconvolved widefield image. DNA is counterstained with DAPI (blue). **d.** H2AK119ub1 nChIP profile for chromosome 3 from iXist-Chr3 mESCs showing H2AK119ub1 gain across the chromosome after 24 h of Xist induction (grey line, two merged replicas^20^ is unaffected in ΔMCPH1 (blue line, two merged replicas). Chromosome sub-compartments depicted in lower bar: A1 dark red, A2 light red, B1 dark blue, B2 light blue, dark grey uncertain compartment assignment. Arrow indicates the location of Xist transgene integration.

Whilst H2AK119ub1 IF clearly indicates enrichment on chromosomes neighbouring chromosome 3 (Fig. 6c), nChIP-seq, which, as noted for parental iXist-Chr3 cells, suggests localisation to B-compartment regions at the distal end of chromosome 3 (Fig. 6d), did not identify Xist-dependent gain of H2AK119ub1 on any other chromosomes. Similarly, analysis of ChrRNA-seq data (Extended Data Fig. 6f) did not identify Xist-dependent gene silencing of regions on other chromosomes.

### Dispersed Xist RNA following HNRNPU depletion retains localisation to Xi A1-sub-compartments

The RNA-binding protein HNRNPU functions in Xist RNA localisation, with HNRNPU depletion resulting in nucleoplasmic dispersal of Xist RNA^11,32–34^. HNRNPU binds directly to Xist RNA^11,33^ and has been suggested to tether Xist molecules (and other target RNAs) to chromatin via the N-terminal SAP and C-terminal RGG domains, which are implicated in DNA/chromatin and RNA binding, respectively. To further analyse HNRNPU function in Xist localisation, we engineered an FKBP12^F36V^ degron tag into HNRNPU in interspecific XX mESCs that have doxycycline-inducible Xist and BglSL/BglG-Halo tagged Xist RNA (Extended Data Fig. 7a, left), followed by engineered deletion of the *Mcph1* gene as described above (Extended Data Fig. 7a, right).

HNRNPU depletion resulted in nucleoplasmic dispersal of Xist RNA in both the presence and absence of MCPH1 (Extended Data Fig. 7b,c), consistent with prior studies^11,32–34^. We used BglSL/BglG-Halo tagging of Xist and super-resolution 3D-SIM imaging to quantify features of the dispersed Xist RNPs on a WT background, revealing an increased volume and reduced density of Xist RNPs, and in addition an increase in the number of Xist molecules (Extended Data Fig. 7d-f). Nuclear fractionation analysis following HNRNPU depletion, again on a WT background, demonstrated that the exclusive association of Xist RNA with chromatin vs nucleoplasmic fractions is markedly reduced (Extended Data Fig. 7g), supporting the proposed function of HNRNPU in tethering Xist RNA to chromatin.

We went on to use ChrRNA-seq to analyse Xist-mediated gene silencing following HNRNPU depletion, both with and without MCPH1 depletion. Unexpectedly we found significant silencing of genes across the X chromosome, albeit to a lesser degree than in WT cells (Fig. 7a). Silencing was more pronounced in the ΔMCPH1 background, in accordance with the modest silencing increase seen in WT ΔMCPH1 mESCs (Fig. 1e). In support of these findings 3D-SIM imaging of Xist RNA FISH on compacted chromosomes in the MCPH1-deficient background, revealed that in addition to dispersed nucleoplasmic signal, localised accumulation of Xist molecules occurs around the Xi chromosome periphery with a single more pronounced cluster likely representing the Xist transcription site (Fig. 7b-e).

**Figure 7.**
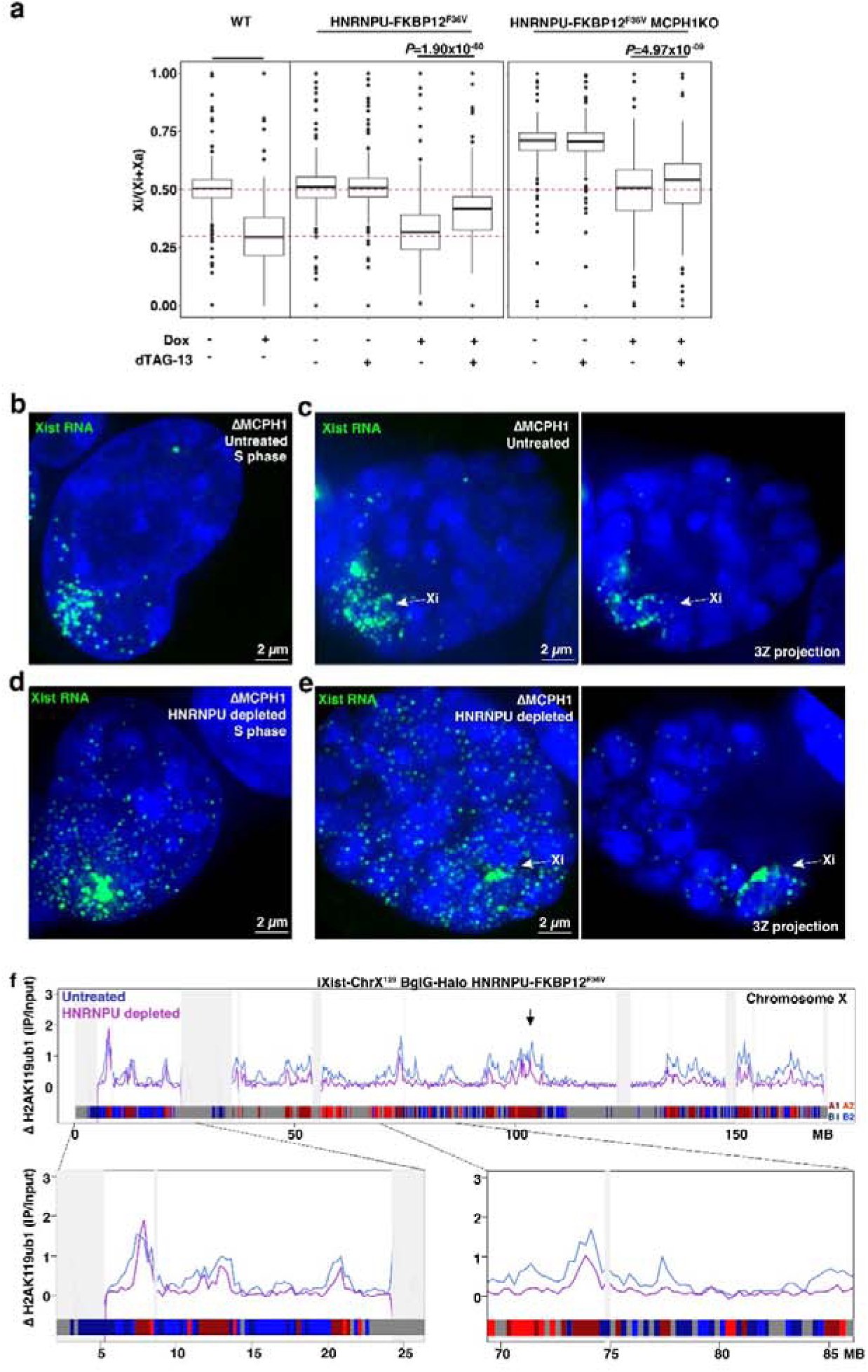
Xist localisation in HNRNPU-depleted ΔMCPH1 mESCs. **a.** ChrRNA-seq in HNRNPU-FKBP12^F36V^ XX mESCs showing silencing of genes on the inactive X chromosome is impaired after depletion of HNRNPU (26 h dTAG-13 treatment, 24 h dox induction of Xist), (n=258 genes, merged experimental duplicates, paired T-test, WT data from^19^, and in ΔMCPH1 cells (n= 248 genes, merged experimental duplicates, paired T-test). **b.** Maximum intensity projection of the deconvolved widefield image of Xist RNA FISH (green) in an untreated ΔMCPH1 HNRNPU-FKBP12^F36V^ cell at S-phase cell (relaxed chromosome morphology), and **c**. an untreated ΔMCPH1 HNRNPU-FKBP12^F36V^ cell with condensed chromosome morphology. Right panel shows 3z maximum intensity projection of the deconvolved widefield image. **d,e.** As for **a,b** after HNRNPU depletion by treatment with dTAG-13. DNA is counterstained with DAPI (blue). **f.** H2AK119ub1 nChIP profile on the X chromosome in HNRNPU-FKBP12^F36V^ mESCs showing enrichment of H2AK119ub1 across the X chromosome after 24 h of Xist induction in the presence (blue line, two merged replicas) and absence of HNRNPU (purple line, two merged replicas), with representative zoomed in regions referenced in the main text. Blocked out panels (light grey) represent regions/genomic bins where the input signal is significantly different (three standard deviations) from its mean value as computed across the X chromosome (see Materials and Methods). Chromosome sub-compartments depicted in lower bar: A1 dark red, A2 light red, B1 dark blue, B2 light blue, dark grey uncertain compartment assignment. The arrow shows the location of the Xist gene.

Localisation of Xist to the periphery of compacted Xi chromosomes in HNRNPU-depleted cells suggests that preferred association with A1-sub-compartments is retained. To test this directly, we analysed Xist-dependent H2AK119ub1 gain across Xi by nChIP-seq. As shown in Fig. 7f, H2AK119ub1 peaks continue to co-localise with A1-sub-compartments in HNRNPU-depleted cells, despite reduced signal across the Xi. We did nevertheless observe differences, with sharper peaks and lower signal at locations immediately flanking major sites (Fig. 7f, expanded viewpoints). Consistent with this observation, a lower H2AK119ub1 signal was associated uniquely with A2-sub-compartments, which are typically juxtaposed with A1-sub-compartments (Fig. 7f, Extended Data Fig. 7h). This distinct profile of H2AK119ub1 gain across the Xi in HNRNPU-depleted cells was unaffected by the depletion of MCPH1 (Extended Data Fig. 7i).

## Discussion

In this study, we show that preferred association sites for Xist RNA on the X chromosome correspond to A1-sub-compartments and that these regions arrange towards the periphery of compacted G1 and G2 chromosomes in ΔMCPH1 cells. We observe the same peripheral pattern of Xist on compacted chromosomes in differentiated C127 cells, indicating that A1-sub-compartments function as the principal sites for Xist localisation during establishment and maintenance of XCI. This observation is in keeping with early work that found Xist localises to gene-rich G-light bands on metaphase chromosomes from differentiated rodent cell lines^35^, and with high-resolution mapping of initial Xist association sites based on mapping of formaldehyde cross-linked chromatin-Xist RNA by high-throughput sequencing analysis^13^. In the SETDB1-depletion model, in which Xist levels are elevated 4-5-fold^15^, analysis of compacted chromosomes showed retention of A1-sub-compartment association and additionally, overspill of Xist molecules onto neighbouring chromosomes. We did not detect accumulation of Xist-mediated chromatin modification (H2AK119ub1) or gene silencing on other chromosomes using molecular assays, and therefore, we cannot directly infer the nature of non-X association sites. This was also the case for the iXist-Chr3 transgene line, in which we observed significant Xist RNA overspill. We interpret these observations as indicating the arrangement of chromosomes neighbouring the Xist-transcribing chromosome is heterogeneous on a cell-to-cell basis, in line with previous reports^36^, and that chromatin modification/gene silencing mediated by spillover Xist is thus distributed at a low-level genome-wide.

Our analysis of XCI in ΔMCPH1 XX mESCs demonstrates that the compacted configuration of chromosomes at G1 and G2 does not impede the spread of Xist RNA from the transcription site to positions along the length of the X chromosome. Indeed, gene silencing progresses somewhat faster in this scenario. Co-localisation of newly synthesised and pre-synthesised Xist molecules on compacted chromosomes, as determined by RNA-SPLIT, further shows that Xist molecules are dynamically translocating despite the relatively compacted chromosome configuration. These findings raise interesting questions about the mechanism by which Xist molecules are transferred along the chromosome. Previous work suggested an important role for 3D chromosome organisation, with Xist association sites correlating with high-frequency contact sites for the Xist locus^13^. A1-sub-compartments have a high frequency of interaction with one another in *cis*, which is consistent with this idea.

However, the 3D arrangement of compacted Xi chromosomes is difficult to reconcile with the proposal that Xist RNA diffuses from the transcription locus to sites that are in proximity due to the 3D-configuration of the chromosome. In a parallel study in which we have developed live-cell tracking of Xist molecules, we conclude that Xist RNPs translocate along the chromosome via contact-mediated transfer (manuscript submitted). Taking this concept into consideration, we propose that interactions between immediately adjacent A1-sub-compartments afford an opportunity for contact-mediated transfer of Xist molecules between successive sites, both in decondensed interphase chromatin and on compacted chromosomes in MCPH1 deficient cells (Fig. 8). Similarly, we propose that in the iXist-Chr3 transgene model where molecular and imaging analyses indicate Xist associates with B-compartment sites on chromosome 3, contact mediated transfer of Xist molecules would occur during interaction of immediately adjacent B-compartment regions, possibly linked to the fact that the Xist transgene is integrated into a B-compartment.

**Figure 8.**
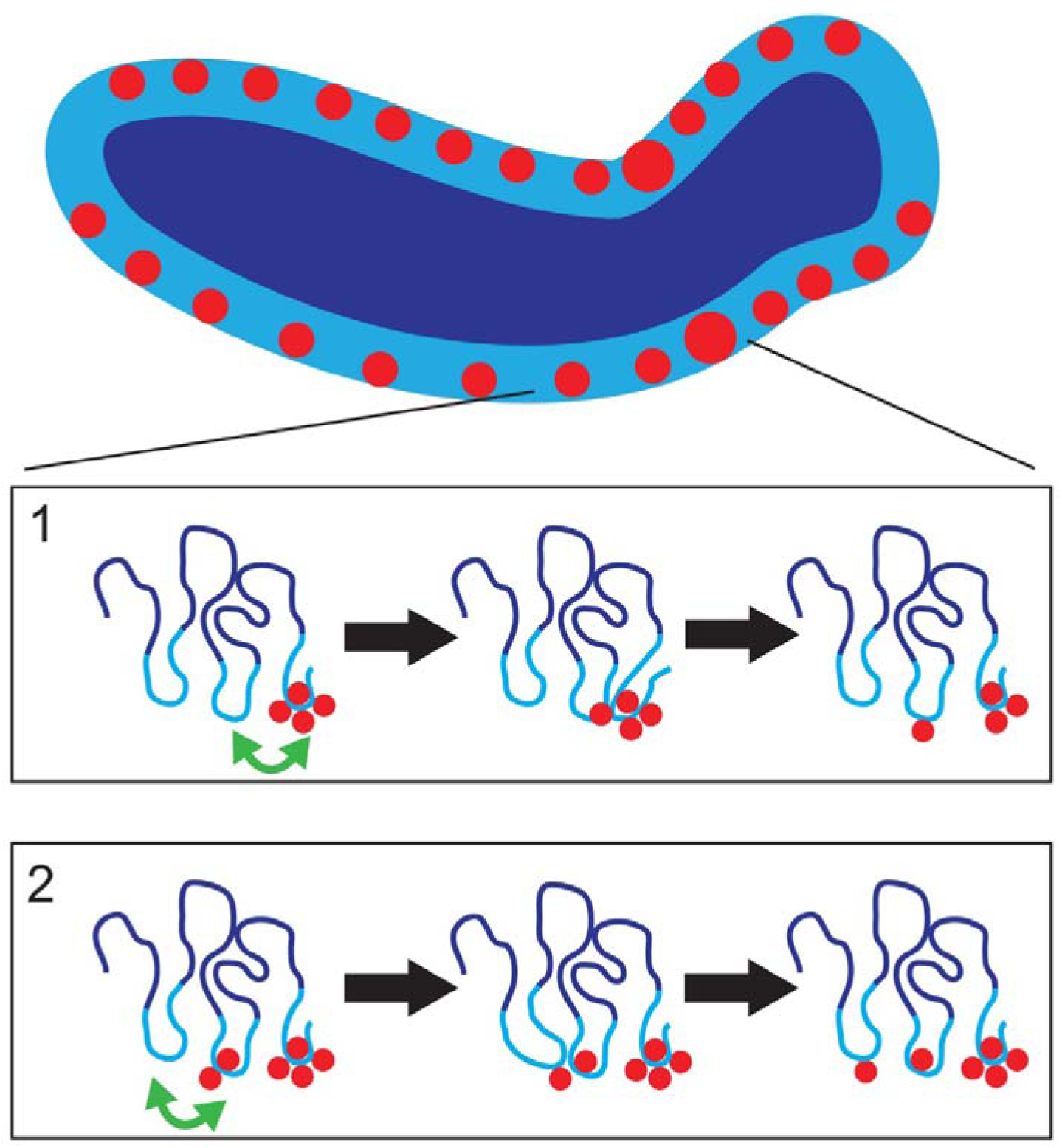
Model for Xist localisation on the Xi in ΔMCPH1 cells. Schematic illustrating organisation of chromosome A-sub-compartments (light blue) and B-sub-compartments (dark blue) on compacted Xi chromosome in G2 ΔMCPH1 cell, indicating Xist RNA at preferred association sites (red circle), and the Xist transcription site (large red circles). Boxes show expanded view indicating condensin II mediated chromosome loops (A-sub-compartments in light blue, B-sub-compartments in dark blue) and a stepwise model for (1) translocation of a single Xist RNA molecule from the Xist transcription site to a neighbouring preferred association site occurring as a result of A1-sub-compartment contact (green arrow), and (2), subsequent translocation of an Xist molecule from the first preferred association site to the next site, again occurring as result of an A1-sub-compartment contact (green arrow).

In the context of the above model, it is interesting that Xist retains a preferred association with A1-sub-compartments in HNRNPU-depleted cells, where localisation is evident across the entire nucleus. Here we envisage that intra-compartment contacts in *cis* determine the preferred localisation of Xist molecules, but that a reduced duration of chromatin association in the absence of HNRNPU results in attenuated spread into A2-sub-compartments, and in molecules being released and diffusing in *trans* to sites across the nucleus. In this view, chromatin anchoring of Xist RNA, facilitated by HNRNPU, and Xist RNA spreading from the transcription site via canonical preferred association sites, are separable processes.

Our analysis of XCI in ΔMCPH1 cells led to unanticipated insights into chromosome organisation at prophase, revealing an inverted inside-outside organisation of A- and B-compartments on compacted chromosomes. We infer that this arrangement reflects chromosome organisation resulting from loading of condensin II but not condensin I, and note that it may explain the constraints that bring about the non-symmetrical condensin II axis observed at prophase^27^. Ongoing transcription is unlikely to be critical for the peripheral localisation of A-compartments, as we observe this arrangement on Xi, where genes are silenced. Instead, we suggest that as condensin II-mediated compaction at prophase releases tethering of heterochromatin from the nuclear envelope, B-compartment interactions in *cis* acting on large chromatin loops predominate, driving the inside-outside configuration of individualised chromosomes. Constraints imposed on loop dynamics by condensin I loading following nuclear envelope breakdown then result in an arrangement that aligns with the linear coordinates of A- and B-compartments, giving rise to the classical G-banding pattern of chromosomes observed at metaphase.

## Materials and Methods

### Cell lines and tissue culture

iXist-ChrX^129^ interspecific *Mus domesticus (129S) x Mus castaneus (Cast)* F1 XX mESC cells with a doxycycline-inducible TetOn promoter driving Xist RNA expression on the *129S* X chromosome^20^, further modified with the introduction of a Bgl stem-loop array in Xist exon 7 and a transgene for expressing BglG-Halo protein integrated into the Rosa26 locus^19^, were used as the parental line from which ΔMCPH1 XX mESCs were initially derived. Targeting via CRISPR-Cas9-mediated homologous recombination was carried out using a plasmid that mediates a 63 bp deletion encompassing most of *Mcph1* exon 2, removing a highly conserved patch leading to loss of protein expression as ascertained by western blot (see below). Briefly, cells were transfected with 2 µg sgRNA (5’-GACGTTTGCAAAGCAACTTG-3’) and 6 µg of homology vector using Lipofectamine 2000 (Life Technologies) according to the manufacturer’s instructions. After 24 h, cells were passaged and treated with 1.75 µg/ml puromycin (Sigma) for a further 48 h. Puromycin selection was then removed, and individual ESC colonies were allowed to grow for ∼10 days, when they were picked and expanded prior to screening by western blot.

iXist-ChrX^Cast^ SETDB1-FKBP12^F36V^ ΔMCPH1 cells were generated by targeting *Mcph1* as described above in XX interspecific mESCs in the SETDB1-FKBP12^F36V^ C7H8 background as detailed previously^15^. Selection in these cells used 3 µg/ml puromycin.

iXist-Chr3 ΔMCPH1 cells were generated by targeting *Mcph1* as described above in XY interspecific *Mus domesticus (129S) x Mus castaneus (Cast)* F1 XY mESC cells that incorporate a multicopy tetracycline inducible Xist transgene randomly integrated on chromosome 3 as detailed previously^20^. Selection in these cells used 3 µg/ml puromycin.

Targeting of iXist-ChrX^129^ interspecific *Mus domesticus (129S) x Mus castaneus (Cast)* F1 XX mESC cells to generate HNRNPU-FKBP12^F36V^ mESCs) were derived by sequential targeting of the N-terminal of the first allele and then the C-terminal of the second allele of the endogenous *Hnrnpu* locus on chromosome 1. Targeting was achieved using CRISPR-Cas9 mediated homologous recombination with a plasmid encoding the 324 bp FKBP12^F36V^ cassette, a 27 bp (i.e 9 amino acid) Gly/Ser linker and 0.5 kb arms of homology. Cells were transfected with 2 µg sgRNA (N-terminal targeting: 5’ - GCCAGCAGTCAGCGGGGAAC-3’ or C-terminal targeting: 5’-AAACAGTCGACTTCTTGTGA-3’) and 2 µg homology vector using Lipofectamine 2000.

Western blotting with an anti-HNRNPU antibody was used to detect the expected increase in protein size and the subsequent loss of the HNRNPU-FKBP12^F36V^ band following 2 h of dTAG-13 treatment. Subsequent retargeting of *Mcph1* in these HNRNPU-FKBP12^F36V^ cells was carried out as detailed above.

mESCs were routinely cultured on gelatinised plates with mitomycin-C (Sigma-Aldrich) inactivated mouse fibroblasts as feeders at 37 °C with 5% CO_2_ in a humid atmosphere in Dulbecco’s modified Eagle’s medium (DMEM, Life Technologies). DMEM was supplemented with 10% fetal calf serum (Seralab), 2 mM L-glutamine, 0.1 mM non-essential amino acids, 50 µM β-mercaptoethanol, penicillin (100 U/ml), streptomycin (100 µg/ml) (all Life Technologies) and 1,000 U/ml LIF (made in-house). As detailed previously, Xist expression was induced by the addition of 1 µg/ml doxycycline (Sigma) for 24 h^20^. mESCs were treated for 2 h with 0.1 µM dTAG-13 (Tocris Bioscience) when relevant to deplete SETDB1 or HNRNPU. In the case of depletion in conjunction with Xist induction, dTAG-13 was applied 2 h prior, and then throughout Xist induction with doxycycline. Where removal of the feeder cells was desirable, mESCs were pre-plated and then grown feeder-free for several passages.

Mouse mammary gland epithelial (C127) cells were grown at 37 °C with 5% CO_2_ in a humid atmosphere in DMEM supplemented with 10% fetal calf serum, 2 mM L-glutamine, 0.1 mM non-essential amino acids, 50 µM β-mercaptoethanol, penicillin (100 U/ml), and streptomycin (100 µg/ml). C127 cells were targeted to delete *Mcph1* exon 2 using the same CRISPR-Cas9-mediated homologous recombination approach described above but selected using 4 µg/ml puromycin. Of note, in targeted C127 cells, *Mcph1^-/-^* was caused by a small insertion/deletion in the same exon 2 patch, likely reflecting the greater propensity for NHEJ rather than HR-mediated CRISPR-Cas9 repair in somatic cells.

### Western blotting

Protein expression was determined by western blotting of nuclear extracts or whole cell lysates. Briefly, for nuclear extracts, cell pellets from pre-plated cells were resuspended 10 volumes of Buffer A (10 mM HEPES pH 7.9, 1.5 mM MgCl_2_, 10 mM KCl, 1 mM DTT, 1 mM PMSF, 1x complete EDTA-free protease inhibitor tablet (Roche), then incubated for 15 min on ice. Cells were then centrifuged at 2,000 g for 5 min at 4 °C, after which the supernatant was discarded. Cell pellets were then resuspended in 3 volumes of Buffer A + 0.1% NP40, incubated on ice for 15 min, and centrifuged at 4,000 g for 5 min at 4 °C, after which the supernatant was discarded. Cell pellets were then resuspended in 1 volume of lysis buffer (20 mM Tris pH 7.5, 200 mM NaCl, 1% Triton X-100, 0.1% sodium deoxycholate, 1 mM DTT, 1 mM PMSF, 1x complete EDTA-free protease inhibitor tablet, 25 U/ml SuperNuclease (Sino Biological) before being passed through a 26G needle 10 times. Samples were then incubated on ice for 1 h, centrifuged at 20,000 g for 30 min at 4 °C, and the supernatant was collected. For whole cell extracts, cell pellets from pre-plated cells were resuspended in RIPA buffer (100 mM Tris pH 7.5, 1 mM EDTA, 0.5 mM EGTA, 1% Triton X-100, 0.1% sodium deoxycholate, 0.1% SDS, 140 mM NaCl, 1x complete EDTA-free protease inhibitor tablet). Samples were separated on a polyacrylamide gel and transferred onto a PVDF membrane using quick transfer. Membranes were blocked for 1Oh at room temperature in 10Oml of TBST (TBS, 0.1% Tween) containing 5% w/v Marvel milk powder, then incubated overnight at 4 °C with the appropriate primary antibody. Blots were then washed four times for 10Omin with TBS-T and incubated for 40Omin with a 1:2000 dilution of a secondary antibody conjugated to horseradish peroxidase (anti-mouse or anti-rabbit IgG HRP (VWR NA931 or NA934, respectively). After washing four times for 5Omin with TBST, bands were visualised using ECL Clarity (BioRad). Some western blots used secondary antibodies raised against rabbit or mouse IgG conjugated to IR680RD or IR800CW dyes (LiCor 926-32210, 926-32211, 926-68070, or 926-68071) 1:15,000 with blots subsequently visualised using an Odyssey Fc western blot imager (LiCor Biosciences). All antibody dilutions were made in blocking buffer. Primary antibodies were as follows: Rabbit anti-MCPH1 (Cell Signalling 4120) 1:1,000, rabbit anti-SETDB1 (Proteintech 11231-1-AP) 1:750, rabbit anti-HNRNPU (produced in-house), 1:200. Mouse anti-TBP (Abcam ab51841) 1:2,000 or rabbit anti-METTL3 (Abcam ab195352) 1:1000 were used as loading controls.

### Standard Xist-FISH

Cells were plated on gelatinised 22 x 22 mm coverslips with inactivated feeders in a 6-well dish for 48 h prior to Xist RNA-FISH. For 26 or 24 h prior to Xist RNA-FISH, 0.1 µM dTAG-13 and/or 1 µg/ml doxycycline was applied as required. Briefly, coverslips were then washed in PBS and fixed using 3.7% formaldehyde in PBS (Sigma) for 10 min at room temperature. After washing in PBS, coverslips were permeabilised with 0.5% Triton X-100 (Sigma) at room temperature for 10 min, then washed again with PBS. Coverslips were then dehydrated in sequential 5 min washes of 80%, 95% and 100% EtOH, briefly dried and then hybridised with 15 µl of fluorescently labelled Xist probe overnight in a dark, humid chamber at 37 °C. Fluorescently labelled Xist FISH probe was generated by nick translation (Abbott Molecular) of a plasmid encoding 15.7Okb mouse Xist cDNA (detailed previously^37^). 50 ng of labelled probe was precipitated per coverslip, with 10Oµg salmon sperm DNA (Invitrogen), 1:10 volume 3OM NaOAc (pH 5.2), and 3 volumes of 100% ethanol. After washing in 70% ethanol, the pellet was dried, resuspended in 6Oµl of deionised formamide (Sigma) per coverslip, and denatured at 75 °C for 7Omin before quenching on ice. The probe was then diluted in 6Oµl 2x hybridisation buffer (5x SSC, 12.5% dextran sulphate, 2.5Omg/ml BSA (NEB), 25 mM VRC (BioLabs; preheated to 65 °C for 5 min). The following day, coverslips were carefully floated off the Parafilm with PBS, returned to a 6-well dish, then washed three times with 2x SSC/50% formamide in PBS, followed by three washes in 2x SSC. All SSC washes were carried out in a water bath at 42 °C. Coverslips were then mounted onto Superfrost Plus microscopy slides (VWR), using Vectashield containing 4,6-diamidino-2-phenylindole (DAPI) (Vector Labs) and sealed with nail varnish. Samples were stored at 4 °C and analysed on an inverted fluorescence Axio Observer Z.1 microscope using a PlanApo ×63/1.4ONA oil-immersion objective. Images were acquired using AxioVision software.

### Xist-FISH and Xist-ImmunoFISH for super-resolution 3D-SIM imaging

Cells were seeded 24 h prior to FISH / ImmunoFISH on No. 1.5H (170 μm ± 5 μm) coverslips (Marienfeld) in a 6-well dish and treated with doxycycline and/or dTAG-13 as appropriate. The following day, coverslips were washed twice in PBS before fixation with 3% formaldehyde in PBS for 10 min at room temperature. Formaldehyde was then replaced stepwise over three washes with PBST (PBS/0.05 % Tween 20), then washed a further two times in PBST prior to permeabilising with 0.5% Triton X-100 in PBS for 10 min at room temperature. Coverslips were then washed twice with PBST before blocking on Parafilm in a humid chamber at room temperature with 3% BSA, 5% normal goat serum, 0.5% fish gelatine in PBST with 30 µl of 40 U/µl RNasin Plus per 600 µl of block, for 15 min at room temperature. For Immuno-FISH, coverslips were placed onto primary antibodies diluted in blocking solution with RNasin on Parafilm in a humid chamber for 1 h at room temperature (mouse anti-phospho-Histone H3 Ser10 clone 3H10 (Merck 05-806, 1:1,000) or rabbit anti-H2AK119ub1 (Cell Signalling 8240S, 1:1,000). Coverslips were then washed three times in PBST before applying a secondary goat anti-mouse IgG Alexa Fluor 568 (Life Technologies A11031) or goat anti-rabbit IgG Alexa Fluor 568 (Life Technologies A11034) diluted 1:1,000 in blocking solution containing RNasin for 30 min at room temperature in a dark, humid chamber. Coverslips were then washed three times with PBST, followed by a further 10 min fixation at room temperature with 3% formaldehyde in PBS. This was exchanged stepwise for PBST and washed a further two times before a final wash in 2x SSC in PBST. The Xist probe was prepared and resuspended in 2x Hybridisation buffer as detailed above, and 10 µl of probe mix was added to coverslips on Parafilm in a dark, humid chamber before overnight incubation at 37 °C. The following day, coverslips were carefully removed by pipetting PBST to gently lift them before placing them in PBST in a 6-well dish. In a fume hood, using pre-warmed solutions and a 42 °C water bath, coverslips were then washed three times in 5% formamide and 2x SSC in PBST for 5 min each wash, then three times in 2x SSC alone in PBST for 5 min each wash. After a PBST wash, coverslips were then incubated with 2 µg/ml DAPI for 10 min at room temperature, then washed with PBS and MilliQ water before being mounted using Vectashield (w/o DAPI) on the reverse (non-frosted) side of a Superfrost microscope slide and sealed with nail varnish.

### Super-resolution 3D-SIM imaging

3D-SIM imaging of fixed samples was initially performed using a DeltaVision OMX V3 system (GE Healthcare) equipped with a 60x/1.42 NA PlanApo oil-immersion objective (Olympus), pco.edge 5.5 sCMOS cameras (PCO), and 405-, 488-, 594-, and 640-nm lasers. Later imaging was performed using a DeltaVision OMX SR system (GE Healthcare; IMSOL) equipped with a 60x/1.5 NA UPLAPO60XOHR oil-immersion objective (Olympus), pco.edge 4.2 sCMOS cameras (PCO), and 405, 488, 568 and 640 nm lasers. Image stacks were acquired at 125 nm z-steps, capturing 15 raw images per z-plane (5 phases, 3 angles). To reduce spherical aberration during image reconstruction, immersion oil with a refractive index of 1.518 was used, or the objective correction collar was set to 0.140, an empirically optimised value for imaging adherent cells (±4 μm). Image reconstruction was performed using softWoRx 7.2.2 (GE Healthcare) with previously obtained, channel-specific optical transfer functions (OTFs)^38^ and a Wiener filter setting of 0.0050. The final reconstructed resolution was approximately 120 nm in the lateral plane (x-y) and ∼320 nm in the axial (z) direction. Quality control was performed using SIMcheck^39^. Correction of chromatic shift between the camera channels was carried out using Chromagnon 3D alignment software^40^, utilising same-day 3D-SIM acquisitions of multicolour ethynyl-2’-deoxyuridine (EdU)-labelled C127 epithelial cells as a colocalisation reference^41^. Images of nuclei presented here were further processed using softWoRx to generate deconvolved widefield images when reconstruction was compromised by excessive out-of-focus blur, particularly for deep cells exhibiting condensed chromosomes. 3D rendering of images used for movies were generated using Imaris software.

### Immunofluorescence staining for 3D-SIM

The day after cells were seeded and induced with doxycycline if appropriate (as described above), coverslips were washed twice in PBS before fixation with 3% formaldehyde in PBS, for 10 min at room temperature. Formaldehyde was then replaced stepwise over three washes with PBST, followed by two further washes in PBST, prior to permeabilising with 0.5% Triton X-100 in PBS for 10 min at room temperature. Coverslips were washed twice with PBST, then blocked on Parafilm in a humid chamber with 3% BSA, 5% normal goat serum and 0.5% fish skin gelatine in PBST for 15 min at room temperature. Coverslips were placed onto primary antibodies diluted in blocking solution on Parafilm in a humid chamber for 1 h at room temperature. Antibodies and concentrations used: mouse anti-phospho-Histone H3 (Ser10) clone 3H10 as above, rabbit anti-H2AK119ub1 as above, rabbit anti-CIZ1 (N-terminal 1794, 1:1,000^12^), rabbit anti-RNA Polymerase II CTD repeat YSPTSPS (phospho S2) (Abcam. ab5095, 1:100), rabbit anti-H3K4me3 (Abcam ab8580, 1:100), mouse anti-H3K27me3 (Active Motif 61017, 1:100). Coverslips were then washed three times in PBST before applying a secondary goat anti-mouse IgG Alexa Fluor 568 (as detailed above), goat anti-rabbit IgG Alexa Fluor 568 (as detailed above), goat anti-mouse IgG Alexa Fluor 488 (Life Technologies, A11029) or goat anti-rabbit IgG Alexa Fluor 488 (Life Technologies, A11008) as appropriate in blocking solution for 30 min at room temperature in a dark, humid chamber. Coverslips were then washed three times with PBST, followed by a further 10 min fixation at room temperature with 3% formaldehyde in PBS. This was exchanged stepwise for PBST, washed twice, then incubated with 2 µg/ml DAPI for 10 min at room temperature, then washed with PBS and MilliQ water before being mounted using Vectashield (w/o DAPI) on the reverse (non-frosted) side of a Superfrost microscope slide, and sealed with nail varnish.

### Xist-Halo staining for 3D-SIM

The day after cells were seeded and induced with doxycycline (as described above), ES media were changed to include 50 nM diAcFAM HaloTag ligand (Promega), and cells were further incubated for 45 min. Cells were then washed by incubating with ES media for 15 min. Doxycycline was maintained throughout. Coverslips were then washed twice with PBS before being fixed in 2% formaldehyde in PBS at room temperature for 10 min before coverslips were washed with PBST, permeabilised with 0.5% Triton X-100 at room temperature for 10 min, washed twice with PBST and incubated with 2 µg/ml DAPI in PBS for 10 min. Coverslips were subsequently washed briefly in PBS, then MilliQ water, before being mounted as described above for 3D-SIM imaging.

### RNA-SPLIT

RNA-SPLIT was carried out as detailed previously^19^. The day after cells were seeded and induced with doxycycline (as described above), ES media were replaced to include 50 nM diAcFAM HaloTag ligand, and cells were incubated for an additional 45 min. Cells were then washed by incubating with ES media for 15 min. For the second pulse of HaloTag ligand, ES media was changed to include 50 nM JF585 HaloTag ligand (kindly provided by Luke Lavis, HHMI Janelia) and incubated for 45 min before washout with ES media for 15 min.

Doxycycline was maintained throughout. Coverslips were then processed as for Xist-Halo imaging as detailed above. The expansion phase refers to analysis after 1.5 h of doxycycline induction, and the steady-state phase to analysis after 24 h of doxycycline induction.

### Chromatin RNA-seq (ChrRNA-seq) and data analysis

Chromatin RNA was extracted from one confluent 15Ocm dish of mESCs, pre-plated, then expanded in advance to remove feeders. Cells were previously treated with 1 µg/ml doxycycline and/or 0.1 µM dTAG-13 as appropriate. Briefly, cells were trypsinised and washed in PBS before being lysed with 1.5 ml RLB (10OmM Tris pH 7.5, 10OmM KCl, 1.5OmM MgCl_2_, and 0.1% NP40) on ice. Centrifugation through a sucrose cushion (24% sucrose in RLB) was used to purify nuclei. Nuclei were then resuspended in NUN1 buffer (20OmM Tris pH 7.5, 75OmM NaCl, 0.5OmM EDTA, 50% glycerol, 1 mM DTT, 1x EDTA-free protease inhibitor tablet (Roche)), before addition of NUN2 buffer (20OmM HEPES pH 7.9, 300OmM NaCl, 7.5OmM MgCl_2_, 0.2OmM EDTA, 1 mM DTT, 1% NP40, 1M urea). After 15 min incubation on ice with occasional shaking, centrifugation at 2,800 g pelleted the insoluble chromatin fraction. The supernatant removed at this point was regarded as soluble nucleoplasmic fraction and saved where appropriate for qRT-PCR analysis. The chromatin pellet was resuspended in TRIzol (Invitrogen) by passing multiple times through a 21G needle. This was used either for qRT-PCR to analyse the nucleoplasmic/chromatin ratio of RNA or further purified for high-throughput sequencing. Chromatin-associated RNA was then purified via TRIzol/chloroform extraction and isopropanol precipitation. Samples were then further purified and concentrated using RNA Clean and Concentrator-5 kit (Zymo Research), including on-column DNAse treatment, before being stored at -80 °C prior to library preparation: Briefly, 1Oµg of RNA was used as input for library preparation using TruSeq stranded total RNA Gold Library Preparation kit (Illumina), and the library was then further quantified by qPCR using KAPA Library Quantification DNA standards and SensiMix SYBR (Bioline, UK). The libraries were pooled and sequenced using either with Illumina NextSeq 500 (2× 81 bp) or Illumina NovaSeq X Plus platform (2× 150 bp). ChrRNA-seq was analysed as follows: The ChrRNA-seq data-mapping pipeline was similar to that in^20^. Briefly, the raw fastq files of read pairs were first aligned to an rRNA build by bowtie2 (v2.3.2)^42^ and rRNA-mapped reads were discarded. The remaining unmapped reads were aligned to the “N-masked” genome (mm10) with STAR (v2.5.2b) using parameters *“–outFilterMultimapNmax 1 –outFilterMismatchNmax 4 –alignEndsType EndToEnd --twopassMode Basic --outSAMstrandField intronMotif --outFilterMultimapNmax 1 --outFilterMismatchNoverReadLmax 0.06”* for all the sequencing libraries^43^. Unique alignments were retained for further analysis. We made use of 23,005,850 SNPs present between *Cast* and *129S* genomes and employed SNPsplit (v0.2.0; Babraham Institute, Cambridge, UK) to split the alignment into distinct alleles (*Cast* and *129S*) using the parameter “--paired”. The allelic read numbers were counted by the program featureCounts^44^ (-t transcript -g gene_id -s 2) against gene annotation, and the alignments were sorted by Samtools (v1.16)^45^. BigWig files were generated by Bedtools (v2.25)^46^ and inspected by IGV^47^. For biallelic analysis of gene expression, counts were normalised to one million mapped read pairs (as CPM) by the edgeR R package. Genes with at least 10 SNP-covering reads across all the samples were further taken to calculate the allelic ratio, i.e. Xi/(Xi+Xa), where Xi (*129S*) and Xa (*Cast*) indicate reads from inactive and active allele, respectively.

The degree of silencing was primarily quantified as the difference in allelic ratios between uninduced and induced samples. Silencing deficiency was compared among different categories of genes, including initial X-linked gene expression level, and promoter chromatin landscape^20^ and gene silencing kinetics^48^. Boxplots were generated using R (v4.2.1) ggplot2 packages.

### Native H2AK119ub1 ChIP-seq

Xist induction by addition of 1.5 µg/ml doxycycline plus/minus 0.1 µM dTAG-13 to pre-plated ESCs as required. Chromatin was made from 50 million ESCs lysed in RSB (10OmM Tris pH 8, 10OmM NaCl, 3OmM MgCl_2_, 0.1 % NP40, 5 mM *N*-ethylmaleimide (NEM, Sigma)).

Nuclei were resuspended in 4Oml of RSB supplemented with 0.25OM sucrose, 3OmM CaCl_2_, protease inhibitor (Roche), and 5 mM NEM, then treated with 200OU of MNase (Fermentas) for 5Omin at 37 °C, inverting tubes every minute. MNase activity was quenched with 8Oµl of 0.5OM EDTA before centrifugation at 5,000Orpm for 5Omin at 4 °C. The supernatant was retained and transferred to a fresh tube whilst the chromatin pellet was incubated, rotating at 4 °C for 1 h in 300Oµl of nucleosome release buffer (10OmM Tris pH 7.5, 10OmM NaCl, 0.2OmM EDTA, 5 mM NEM protease inhibitor). The chromatin fraction was then passed five times through a 27G needle before centrifugation at 5,000Orpm for 5Omin. The supernatant was combined with the previously isolated fraction to give the final chromatin sample, which was snap-frozen and used as required for ChIP experiments (after the size of the chromatin fragments was confirmed by agarose gel electrophoresis to be predominantly mono-nucleosomal). For each ChIP reaction, 100Oµl of chromatin was diluted in incubation buffer (10OmM Tris pH 7.5, 70OmM NaCl, 2OmM MgCl_2_, 2OmM EDTA, 0.1% Triton X-100) to 1Oml and incubated with anti-H2AK119ub1 antibody (Cell Signalling 8240S) overnight at 4 °C. All buffers were supplemented with 10OmM NEM. Samples were incubated for 1Oh at 4 °C with 45Oµl protein A agarose beads pre-blocked in incubation buffer supplemented with 1Omg/ml BSA and 1Omg/ml yeast tRNA, then washed five times in ice cold wash buffer (20OmM Tris pH 7.5, 2OmM EDTA, 125OmM NaCl, 0.1% Triton X-100) and once in ice cold TE. DNA was eluted by shaking at 25 °C for 15 min with 1% SDS and 100OmM NaHCO_3_, eluate removed, the elution step repeated and pooled with the first eluate. This, along with an input sample was then purified using the ChIP DNA Clean and Concentrator kit (Zymo Research). Approximately 40Ong of ChIPed DNA was used for library prep with the NEBNext Ultra II DNA Library Prep Kit with NEBNext Single indices (E7645), and the library was then quantified by qPCR using KAPA Library Quantification DNA standards and SensiMix SYBR (Bioline, UK). The libraries were pooled and sequenced using either with Illumina NextSeq 500 (2× 81 bp) or Illumina NovaSeq X Plus platform (2× 150 bp).

### Data analysis of H2AK119ub1 ChIP-seq and RAP-seq

Paired-end raw reads were aligned using Bowtie2 (v2.3.2)^42^ to the mouse reference genome mm10, with key parameters “*--sensitive --no-mixed --nodiscordant --maxins=1000*”. Reads with mapping quality >=7 were retained (-q 7), and all reads flagged by Bowtie2 with a secondary alignment flag were discarded. Reads were then deduplicated using picard-tools (http://broadinstitute.github.io/picard/). The SNPsplit genome preparation was run for the dual-hybrid strain, *129S/Cast*, and allele-specific analysis was performed, assigning each read to either the *Cast* or *129S* reference using the mm10 reference genome masked with the ambiguity nucleobase ‘N’ (N-masking). Profiles for the active and inactive X chromosome were obtained accordingly. Normalisation and binning were implemented using our in-house Python scripts. Per-million read normalisation was applied to both RAP-seq and ChIP data and their corresponding input libraries, and finally, coarse-grained profiles at either 50 kb or 250 kb resolution were calculated.

### Data analysis of gene density and transposable elements

"Reference collections of human and rodent repetitive elements" (http://www.girinst.org/) was used to enumerate SINE and LINE transposable elements, as inferred from the mouse reference genome, mm10. Alternatively, in the analysis shown in Extended Data Figure 6a, SINE repeat element annotations for mm10 were obtained from the UCSC RepeatMasker table (rmsk.txt.gz, April 2021), which is derived from Repbase repeat libraries and is consistent with the above annotation source. GenomeRange^49^ was used to convert transposable element information into an R data frame (https://www.R-project.org/) and to perform downstream analysis with Karyoplote^50^. In particular, the kpPlotDensity function from KaryoploteR was used to generate normalised densities of SINE and LINE transposable elements, shown alongside the RAP-seq and ChIP-seq experiments. A similar approach was used to generate gene density profiles, in which the lists of protein-coding genes and long non-coding genes were extracted from the GENCODE annotation^51^ and passed to GenomeRanges and KaryoploteR for downstream gene density analysis. Both transposable elements and gene density profiles were then binned at either 50 kb or 250 kb resolution for analysis. Boxplots were generated by computing the relative contribution to each compartment/sub-compartment of the normalised signal (IP/Input), both in the WT and in the HNRNPU-FKBP12^F36V^ dataset. Pearson correlation analysis was performed using the ‘cor’ function in R, with subsetting to genomic locations/bins that were not masked (shaded grey areas). Bins were masked when the value of the Input signal was three standard deviations away from the mean, as computed across the entire X chromosome. When two replicates were combined, the masking procedure was applied to both Input signals separately, and the final blacklist was obtained by concatenating the individual replica blacklists. Heatmap data shows the centromeric region of the X chromosome [chrX:5Mb-25Mb obtained from <u>SRR6822773</u>. This corresponds to ESC, Day0, unsplit data. Sub-compartments were computed as in^52^.

### Analysis of chromatin compartments

To compare chromosome compartments with ChIP-seq and RAP-seq enrichment profiles, raw unsplit Hi-C data from undifferentiated mESCs were obtained from (<u>SRR6822773</u>^53^) and processed using ICeCAP^54^, with the capture-specific, oligo-bait flag option off (as non-capture bulk Hi-C data). Briefly, reads were first flashed using FLASH^55^. Both extended reads and unflashed reads were in silico digested at a modified restriction site and paired into chimeric ditags, following the filtering principles detailed in HiCUP^56^ and NGseqBasic^57^. Hi-C paired-end reads were aligned to the N-masked mm10 reference genome as single-end reads using Bowtie2 (v2.3.2)^42^. Allele-specific analysis was implemented using SNPsplit^58^. Read pairs were assigned to 50 kb genomic bins, and a Hi-C contact matrix was generated with Cooler^59^ and Cooltools^60^. Unmappable regions were defined as bins with low read coverage (the bottom 25% of bins) and were excluded from subsequent analysis. Hi-C contact matrices were normalised using Cooler^57^ using the “--cis-only” flag option on individual chromosomes^61^. A-compartments and B-compartments were distinguished based on the sign of the first eigenvalue component, respectively (and gene density analysis). Sub-compartment analysis was performed using the Pearson correlation matrix of the observed Hi-C over expected values, applying KMean analysis to the first eigenvalue, using a method outlined previously^52^. BedGraph files for each chromosome were generated at a 50 kb resolution, with bins assigned to either the A-compartment (identifier value 1) or the B-compartment (identifier value 2), or to sub-compartments A1, A2, B1, B2. Further processing of these files in R was performed to generate diagrams depicting the genomic locations of compartments together with binned ChIP and RAP-seq data.

### C127 RNA-seq

WT and MCPH1 KO C127 cells were washed with PBS and then harvested with a cell scraper. Total RNA was extracted using TRIzol (as per manufacturer’s instructions), then further purified using a RNeasy Mini kit (Qiagen) with on-column DNase treatment (as per manufacturer’s instructions). Each sample was repeated in triplicate. Libraries were prepared using the TruSeq stranded total RNA Gold Library Preparation kit and sequenced as detailed above. Differential gene expression analysis was carried out as follows: in order to call differential gene expression between WT and ΔMCPH1 C127 cells, raw FASTQ files containing paired-end reads were first aligned to a custom rRNA reference using Bowtie2 (v2.3.5.1)^42^ to remove rRNA contaminants. Reads mapping to rRNA were discarded, and the remaining unmapped read pairs were subsequently aligned to the mouse genome (mm10) using STAR (v2.7.9a)^43^. STAR was run with the following parameters: *“--twopassMode Basic --outSAMstrandField intronMotif -- outFilterMismatchNoverReadLmax 0.06 --outFilterMultimapNmax 100–alignEndsType EndToEnd –winAnchorMultimapNmax 100*”. Alignments were sorted and strand split using SAMtools (v1.16)^45^. Differential expression analysis of genes and transposable elements was performed using TEtranscripts (v2.2.1)^62^, applying an adjusted p-value threshold of 0.05 to define significance.

### RASER-FISH and Immuno-RASER FISH

RASER-FISH / Immuno-RASER FISH was developed by adapting a previously published protocol^25^. Briefly, cells were plated at low density on gelatinised 22 x 22 mm No. 1.5H precision coverslips (Marienfeld Superior) in 6-well plates with inactivated feeders overnight prior to the experiment with BrdU/BrdC mix (ratio 3:1) at a final concentration of 10 µM. The next day, coverslips were washed with PBS, then fixed with 4% formaldehyde in PBS at room temperature for 10 min. Coverslips were then rinsed a further three times in PBST before being permeabilised with 0.5% Triton X-100 in PBS at room temperature for 10 min. For RASER Immuno-FISH, after a further three PBST washes, coverslips were blocked in 3% BSA and 5% goat serum in PBS at room temperature for 15 min, before coverslips were inverted onto 100 µl of primary antibody on Parafilm in a humid container at room temperature for 30 min (Rabbit anti-CIZ1, N-terminal 1794, 1:1,000^12^). Coverslips were then washed three times in PBST and incubated with 100 µl secondary antibody (Goat anti-rabbit IgG Alexa Fluor 647, Life Technologies A21245, 1:1,000) on Parafilm in a humid container at room temperature for 30 min. All antibody dilutions were made in block solution. After washing 3 times in PBST coverslips, free aldehydes were quenched by a 10 min room temperature incubation with ammonium chloride (0.13g NH_4_Cl in 50 ml PBS), washed three times with PBST, then treated with RNase A (0.1 mg/ml RNaseA (Sigma) in PBS) and incubated on parafilm in a floating humid chamber at 37 °C for 45 min. Cells were then sensitised for RASER-FISH by treating with 0.5 µg/ml DAPI in PBS (1:10,000) at room temperature for 15 min. After a brief rinse, the PBS volume was reduced to 1 ml in a 6-well dish. The 254 nm UV crosslinker was prewarmed with an ‘auto crosslink’ run, and then cells were exposed to 254 nm UV light for 17 min with the lid off. Coverslips were then inverted onto 60 µl Exonuclease III diluted in 1x Exonuclease III buffer and spotted onto Parafilm (NEB MO206L) in a humid container. This was then incubated at 37°C for 15 min, floating in a water bath. Coverslips were washed 3 times with PBS, then inverted onto 12 µl of probe mix and hybridised overnight in a humid chamber at 37°C. The L1 probe was generated by PCR amplification from iXist-ChrX^129^ ES cell gDNA using the forward primer 5’-GCCTCAGAACTGAACAAAGA-3’ and the reverse primer 5’-GCTCATAATGTTGTTCCACCT-3’, yielding a 1041 bp PCR product that was TA cloned into the TOPO pCR2.1 cloning vector (ThermoFisher Scientific) as per the manufacturer’s instructions and Sanger sequenced to confirm correct amplification of the LINE1 fragment. This L1-containing plasmid was then used as a template from which working stocks of labelled probe were synthesised by PCR amplifying the sequence with the same L1 primers in the presence of 1 mM dUTP-Spectrum Green. Amplification was confirmed by agarose gel electrophoresis, and the product was purified using a DNA Clean and Concentrate kit (Zymo) before elution in 25 μl of pre-warmed elution buffer. The B1 probe was synthesised commercially to include 5’ Alexa Fluor 594 (Invitrogen Life Sciences). The probe was a 1:1 mix of two HPLC-purified 30-base-oligonucleotides: mouse B1-1 5’-gcctggtctacagagtgagttccaggacag-3’ and mouse B1-2 5’-cagcacttgggaggcagaggcaggcggatt-3’ ^63^. The oligonucleotides were resuspended in 1x RASER-FISH hybridisation buffer (2.5x SSC and 6.25% dextran sulphate, 50% formamide in H_2_O) to 100 ng/μl and frozen in single-use aliquots. To prepare the L1 and B1 probe mix for use, 100 ng of L1 probe per coverslip was precipitated as detailed above for the Xist probe. This was then resuspended in 5.5 µl formamide per coverslip, incubated shaking at 42 °C for 30 min, then added to 5.5 µl of 2x RASER-FISH hybridisation buffer (5x SSC, 12.5% dextran sulphate in H_2_O) per coverslip and mixed well. 1 µl (100 ng) B1 probe per coverslip was then added to the mix and the probe mix was denatured at 95 °C for 10 min, before being placed immediately on ice. The following day, coverslips were carefully floated off and placed, cell-side up, into 4x SSC/PBS in a 6-well plate. Coverslips were then washed twice in 2X SSC/PBS at 37 °C for 30 min each in a 6-well dish floating on a water bath, washed once in 1X SSC/PBS at room temperature for 30 min, then rinsed once briefly with MilliQ water before mounting on slides using Vectashield containing DAPI and sealed with clear nail varnish.

### Hi-C chromatin conformation capturing analyses

Hi-C was performed as described^64^ with minor modifications. Cells were dissociated with trypsin and fixed in aliquots of 5x10^6^ cells in media with 1% formaldehyde for 10 min at room temperature, before quenching with 128 mM glycine. Cell pellets were then fixed with 300 mM disuccinimidyl glutarate (DSG) in PBS for 40 min before quenching with 400 mM glycine. Samples were lysed and Dounce homogenised, before nuclei were washed and resuspended in DpnII buffer (NEB). 1% SDS was added and samples were incubated at 65°C for 10 min and quenched with 1% Triton X-100. Subsequently, 400 U of DpnII (NEB) was added and samples were incubated for 10 h at 37°C with interval shaking (1400 rpm, Eppendorf Thermomixer). DpnII was heat-inactivated 65°C for 10 min and DNA ends were biotinylated before being ligated with 50 U of T4 DNA ligase (ThermoFisher, EL0013) in 1x ligase buffer (ThermoFisher, B69), with 1% Triton X-100, and 100 µg/ml recombinant albumin overnight at 16°C with interval shaking. Cross-linking was reversed with 400 µg/mL proteinase K at 65°C overnight with interval shaking. DNA was purified using the DNeasy Blood & Tissue kit (Qiagen, 69504). To assess digestion and ligation efficiency, samples were taken pre-digestion, post-digestion, and after ligation and analysed with the Agilent TapeStation (genomic DNA screen tape). 10 µg of ligated DNA was taken forward per sample. Biotin was removed from unligated ends by T4 DNA polymerase, and libraries sonicated to 300 bp using a Covaris ME220 at 175 W, 20% duty cycle, and 200 cycles per burst for 140 sec. Sonicated DNA was purified with AMPure XP beads. DNA was not size-selected but instead directly pulled down with 10 µl of streptavidin beads per sample. Libraries were then constructed using the NEB ultra II library preparation kit (E7645S) according to the manufacturer’s instructions. Afterwards, the beads were washed 2x in Tween-20 wash buffer, 2x in binding buffer, 1x in MilliQ water, and finally resuspended in 20 µl MilliQ water. 15 µl of each library was PCR amplified and indexed following the NEB Ultra II library preparation kit instructions. The final libraries were purified using AMPure XP beads, pooled and analysed with Illumina 150 bp paired-end sequencing. Sequencing data were pre-processed using the Juicer (v1.6) pipeline^65^. Topologically associated domain (TAD) analysis and Hi-C matrix visualisation was performed using HiCExplorer (v3.7.2) tools^66^. A/B compartmentalisation was analysed and plotted using Cooltools (v0.7.1)^60^.

### Cell fractionation and qRT-PCR

Fractionation for qRT-PCR followed broadly the same protocol used for isolation of ChrRNA detailed below, with the nucleoplasmic fraction being the supernatant removed from the chromatin pellet. Samples for fractionation were taken from a T145 dish of cells pre-plated to remove feeder cells and treated with 0.1 µM dTAG-13 for 2 h prior to the addition of 1 µg/ml doxycycline, meaning that cells were treated for a total of 22 h dTAG-13 and 20 h with doxycycline. RNA was extracted from chromatin and nucleoplasmic fractions using Trizol, then further purified using the Direct-zol RNA miniprep kit with on-column DNase treatment (Zymo Research). cDNA was synthesised from 2 µg of total RNA using a mix of random hexamers, 40 units of RNasin Plus (Promega) in duplicate, one with SuperScript III Reverse Transcriptase as per the manufacturer’s instructions (ThermoFisher) and the other with H_2_0 as a control. qRT-PCR reactions were run on a Rotor-Gene Q (Qiagen) thermocycler with reaction parameters of: 95 °C for 3 min; 40 cycles of 95 °C for 20 s, 60 °C for 20 s; melt curve 55–90 °C. Ct values were calculated after applying a fluorescence threshold of 0.3 such that all samples were in the log amplification phase. Each sample was run in triplicate to obtain an average Ct value that was normalised to Actin. The presented data represent the SEMs of three experiments, including two biological replicates. 1 µl of cDNA was used per qRT-PCR reaction along with SensiMix SYBR (Bioline), 0.3 µM of forward and reverse primers (Xist BM33 5’ TCCAGGCAATCCTTCTTCTTG 3’, Xist BM34 5’ GATCCTGCTTGAACTACTGCT 3’ or Actin BM31 5’ TGTTACCAACTGGGACGACA 3’, Actin BM32 5’ CCATCACAATGCCTGTGGTA 3’). Controls containing no DNA and samples in which no RT enzyme was included in the cDNA reaction were used as negative controls for the PCR reaction.

## Data Availability

The datasets produced in this study are available in the following database: ChrRNA-seq: ENA; Project PRJEB101053 (https://www.ebi.ac.uk/ena). nChIP-seq: ENA; Project PRJEB101053 (https://www.ebi.ac.uk/ena).

## Acknowledgements

We would like to thank Martin Houlard for advice and the CRISPR-Cas9 targeting strategy. Imaging was carried out at the Micron Oxford Advanced Bioimaging Unit at the Department of Biochemistry (University of Oxford), supported by a Wellcome Trust Strategic Award (091911 and 107457/Z/15/Z). This work was supported by Wellcome Trust (215513/Z/19/Z to N.B.) and UKRI (EP/Y029062/1 to N.B.). J.D. is supported by the Lister Institute, the MRC Molecular Haematology Unit (MC_UU_00029/04) and Wellcome (225220/Z/22/Z). G.L. is supported by the BBSRC (BB/T008784/1).

---

**Extended Data Figure 1.**
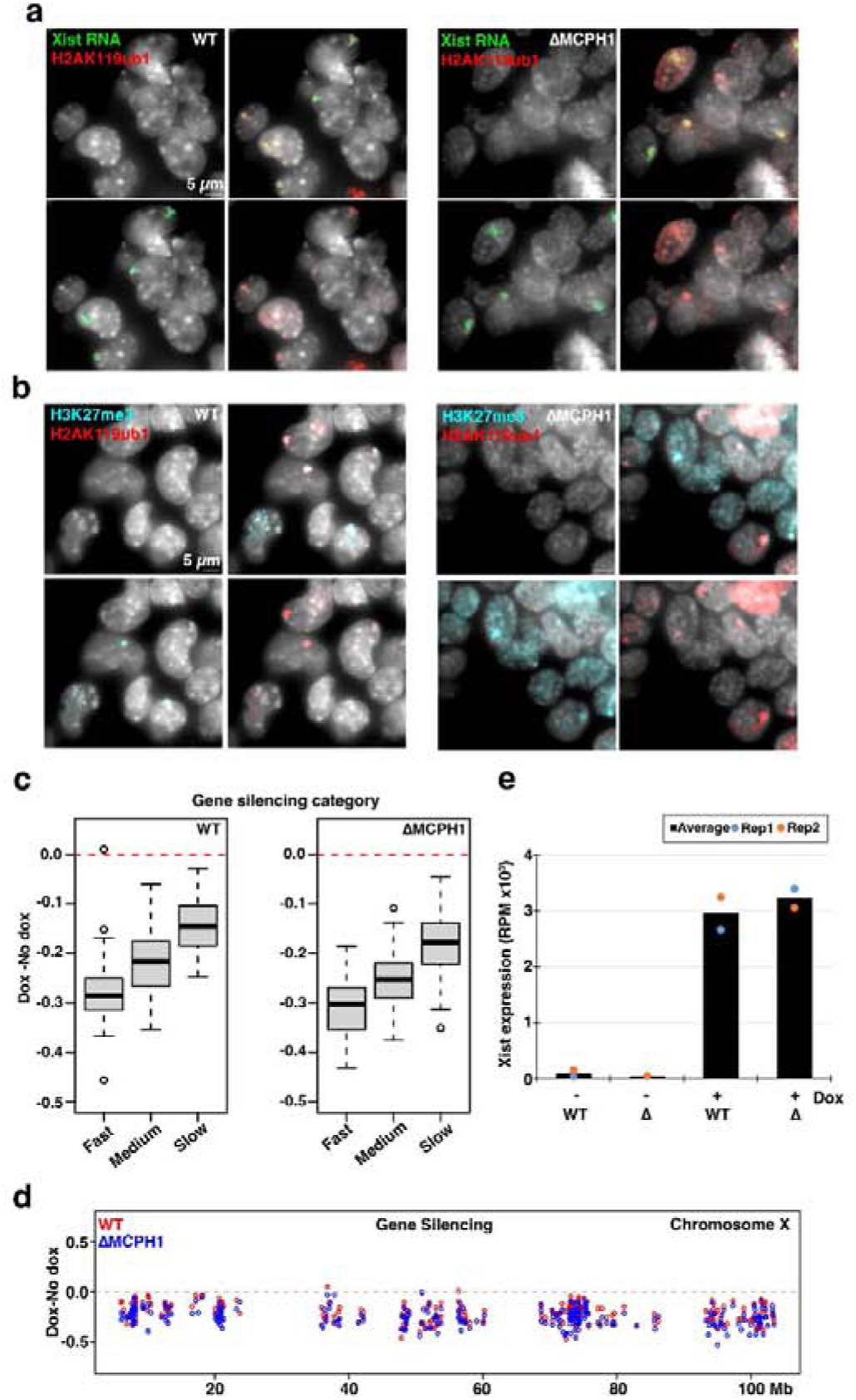
Generation of ΔMCPH1 XX mESCs. **a.** Immuno-FISH of H2AK119ub1 and Xist RNA in WT and ΔMCPH1 iXist-ChrX^129^ BglG-Halo mESCs. **b.** IF for H2AK119ub1 and H3K27me3 in WT and ΔMCPH1 iXist-ChrX^129^ BglG-Halo mESCs. **c.** Analysis of ChrRNA-seq in iXist-ChrX^129^ BglG-Halo mESCs shows no effect of ΔMCPH1 on the Xi silencing dynamics in categories of Xi genes grouped by gene silencing kinetics (fast, medium and slow). All data is from two merged replica experiments. WT from^19^. **d.** Analysis of ChrRNA-seq in iXist-ChrX^129^ BglG-Halo mESCs showing gene silencing occurs across the Xi in ΔMCPH1 (2 merged replica experiments, WT from^19^. **e.** Xist expression levels determined from ChrRNA-seq in iXist-ChrX^129^ BglG-Halo mESCs in the presence^19^ and absence of MCPH1 and +/- dox induction (average of two replicas shown).

**Extended Data Figure 2.**
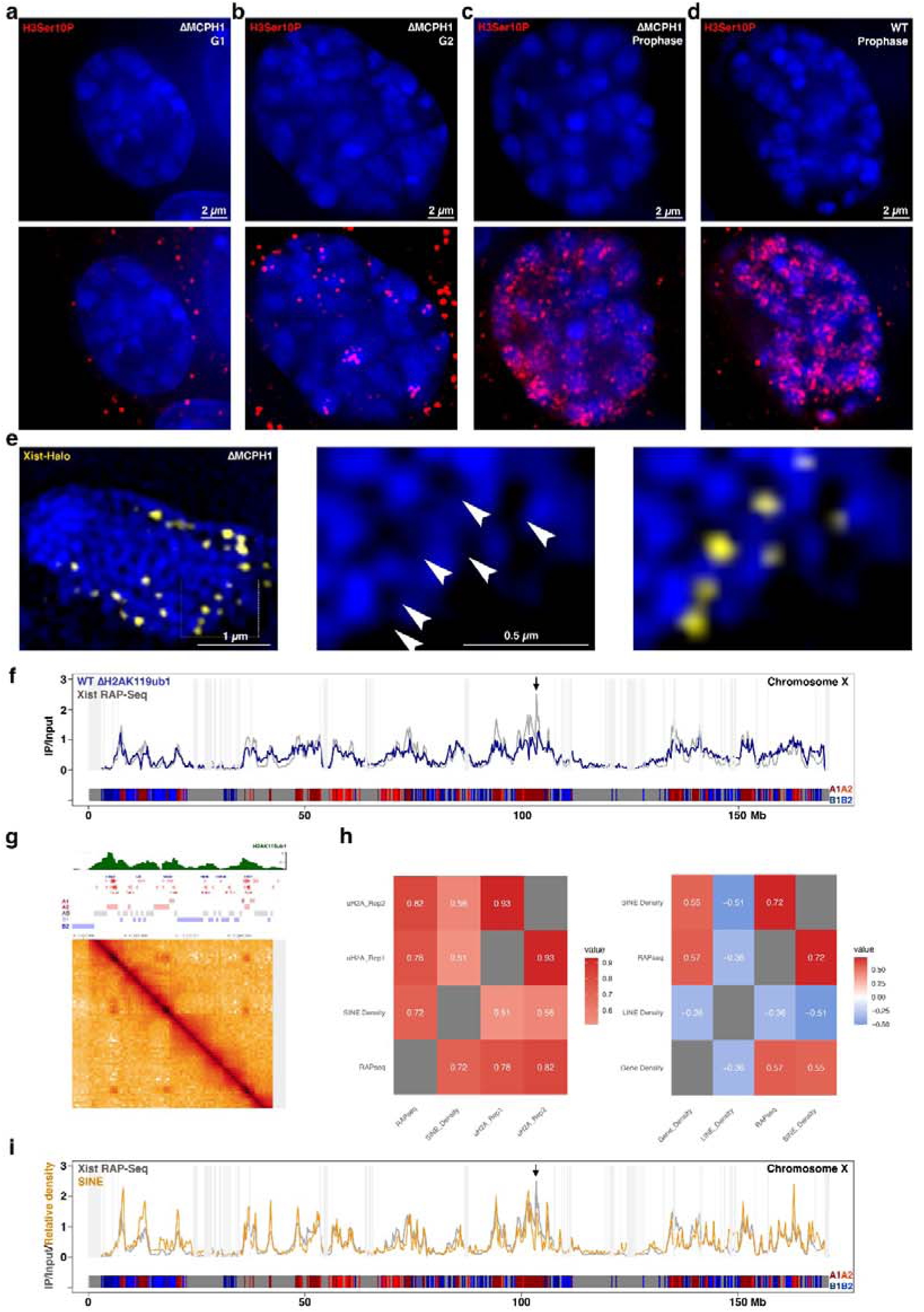
Analysis of XCI in ΔMCPH1 XX mESCs. **a-d.** Deconvolved widefield image of H3Ser10P immunofluorescence in iXist-ChrX^129^ BglG-Halo mESC. A single z from the maximum intensity projection in Figure 2 is shown here for clarity; **a.** ΔMCPH1 G1 cell. **b.** ΔMCPH1 G2 cell. **c.** ΔMCPH1 prophase cell. **d**. WT prophase cell. **e.** Xist-Halo diAcFAM staining of compacted Xi from ΔMCPH1 iXist-ChrX^129^ BglG-Halo mESC. Area shown in zoomed images (right) is indicated with white rectangle. Localisation of Xist foci within interchromatin channels is highlighted with white arrowheads. Images are maximum intensity projections of 3 3D-SIM z-sections. In **a-e** DNA was counterstained with DAPI (blue). **f.** H2AK119ub1 nChIP (blue line) profile in WT iXist-ChrX^129^ BglG-Halo mESCs after Xist induction for 24 h (two merged replicas). The profile is shown relative to Xist RAP-seq data at 48 h^22^ (grey line), and chromosome sub-compartments (A1 dark red, A2 light red, B1 light blue, B2 dark blue, with grey bins representing regions that are masked by the iterative correction where a weight cannot be assigned for normalization) indicated in bar. **g** Heatmap showing Hi-C data from^53^ illustrating enhanced chromatin contact frequency between A1-sub-compartments within indicated region of the X chromosome. Genes are annotated in blue and red. H2AK119ub1 track (green) is from iXist-ChrX^129^ BglG-Halo mESC cells after 24h Xist induction (pooled from 2 replica experiments). Sub-compartments shown in bar are as in **f**. **h.** Pearson correlation of H2AK119ub1 replica ChIP experiments from iXist-ChrX^129^ BglG-Halo mESC cells after 24 h of Xist induction, with 48 h Xist RAP-seq and SINE density (left panel). Pearson correlations for Xi gain of H2AK119ub1, 48 h Xist RAP-seq, SINE density gene density, and LINE1 density across the X chromosome. **i**. Xist RAP-seq (48 h, grey line) and SINE density (ochre line) correlate with A-compartments and anti-correlate with B-compartments, depicted in the lower bar. Compartment annotation: A1 dark red, A2 light red, B1 dark blue, B2 light blue, AB, dark grey compartment assignment not conclusive.

**Extended Data Figure 3.**
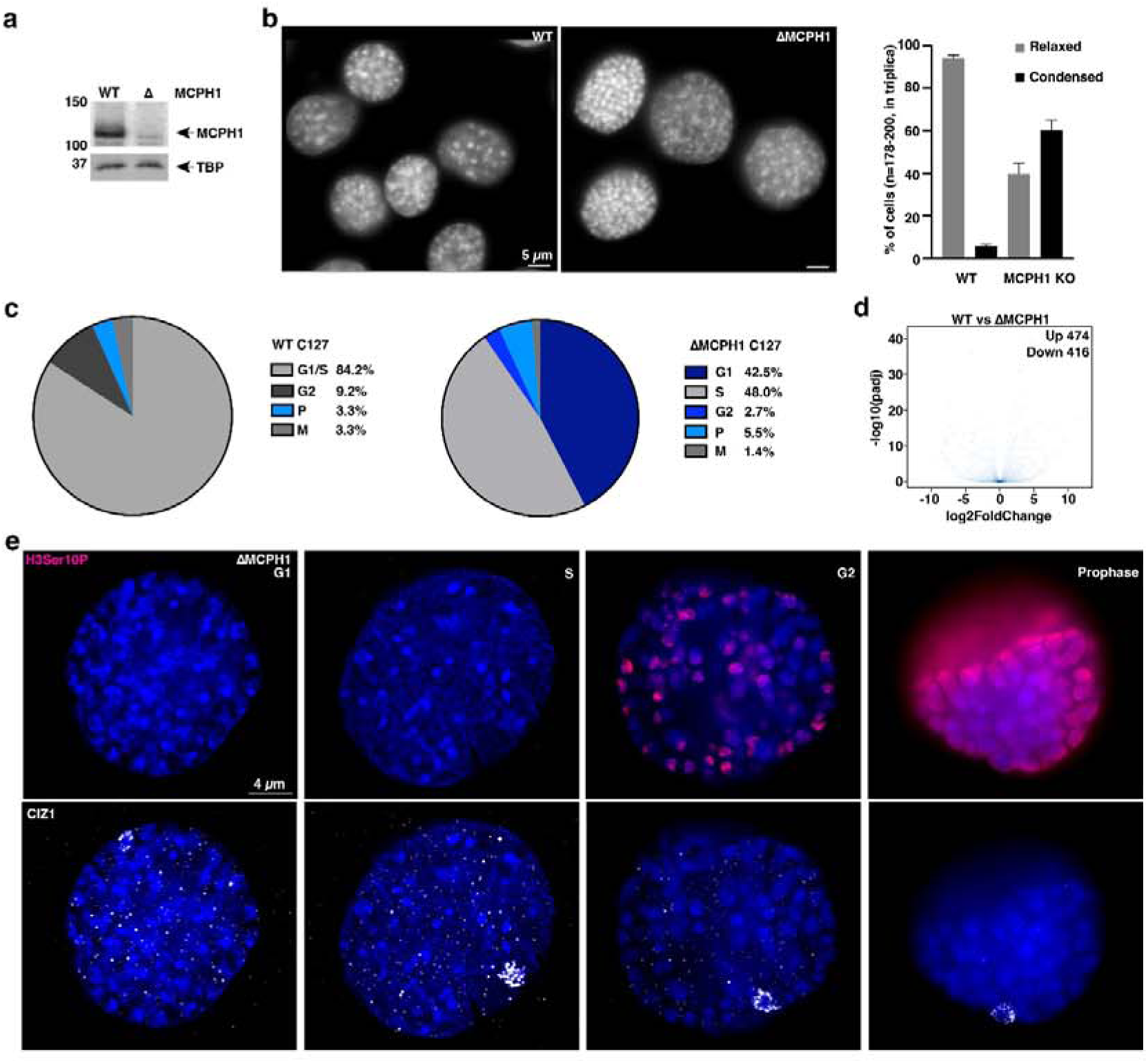
Analysis of ΔMCPH1 C127 XX somatic cells. **a.** Western blot of nuclear extracts from WT and ΔMCPH1 C127 cells probed with anti-MCPH1 (anti-TBP loading control), showing loss of MCPH1 in ΔMCPH1 cells. **b.** Example widefield images of chromosome morphology in C127 WT and ΔMCPH1 cells (DNA counterstained with DAPI), with a histogram (right) showing the scoring of cells with relaxed vs condensed chromatin states. **c.** Pie charts showing cell cycle distribution for WT and ΔMCPH1 C127 cells, defined by H3Ser10P IF and chromosome morphology visualised by DAPI staining of DNA (WT n= 120, MCPH1 KO n=73). **d.** Total RNA-seq of WT and ΔMCPH1 C127 cells (merged triplicate experiments) demonstrating a minimal effect on gene expression (474 genes significantly upregulated, 416 genes significantly downregulated). **e.** H3Ser10P and CIZ1 IF in ΔMCPH1 G1, S, G2 and prophase cells, respectively. DNA counterstained with DAPI (blue). All panels show maximum intensity projections of the deconvolved widefield image.

**Extended Data Figure 4.**
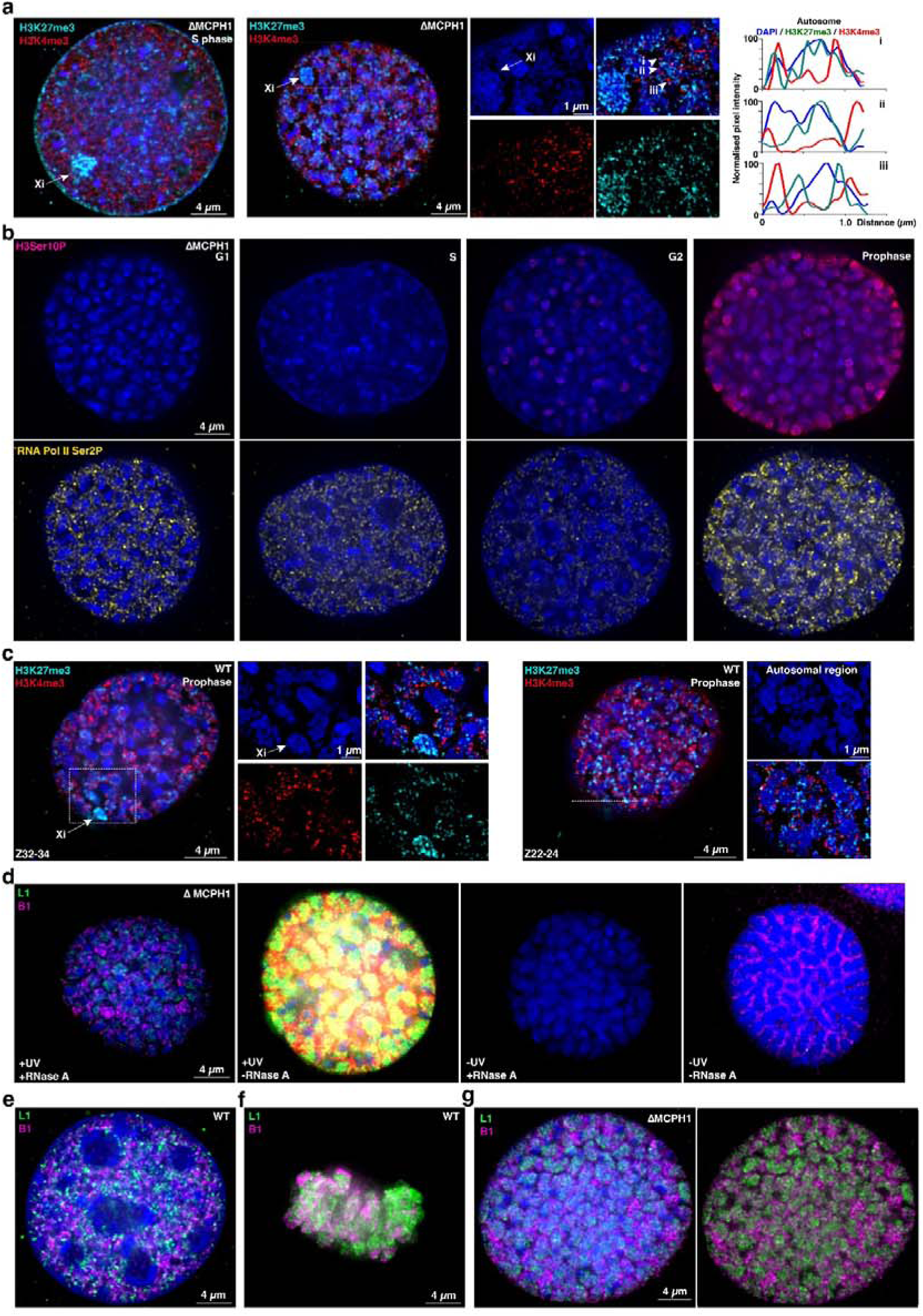
Analysis of XCI and chromosome organisation in ΔMCPH1 C127 XX somatic cells. **a.** IF for H3K27me3 and H3K4me3 in a C127 ΔMCPH1 cell in S-phase with relaxed chromosome morphology (left panel) and a C127 ΔMCPH1 cell exhibiting condensed chromosome morphology (centre panel and insets showing colour split of zoomed area highlighted with white rectangle). H3K27me3 and H3K4me3 distributions across an autosome are depicted IF signal intensity along lines denoted i-iii, with the start of each line indicated with an arrowhead (right panel). **b**. Images showing IF for RNA Pol II Ser2P in ΔMCPH1 C127 cells. Different cell cycle stages were determined from H3Ser10P IF and DAPI counterstaining. **c**. Maximum intensity projections of three z sections of deconvolved widefield images illustrating IF for H3K27me3 and H3K4me3 in WT prophase C127 cells (cell cycle phase determined by cell morphology). Panels and insets of zoomed area in white rectangle highlight staining around Xi (left) and non-Xi (right) regions. **d**. Images of RASER-FISH controls demonstrating UV dependency of RASER-FISH signal (compare first and third panels). RNase A treatment removed the signal from probes binding to RNA (second and fourth panels). **e-f.** Images of RASER-FISH showing L1 and B1 DNA elements in WT C127 cell **e.** at interphase, and **f.** in mitosis, after nuclear membrane breakdown (showing classic banded chromosome pattern with both condensin I and II loaded). **g**. RASER-FISH image showing inside-outside distribution of L1 and B1 DNA elements in a ΔMCPH1 C127 cell with condensed chromosomes. For all panels a white rectangle indicates the region of the zoom view shown in insets. Images of nuclei are maximum intensity projections of deconvolved widefield images unless otherwise stated, with inset zoom images showing a maximum intensity projection of 3 3D-SIM z-sections. DNA is counterstained with DAPI (blue).

**Extended Data Figure 5.**
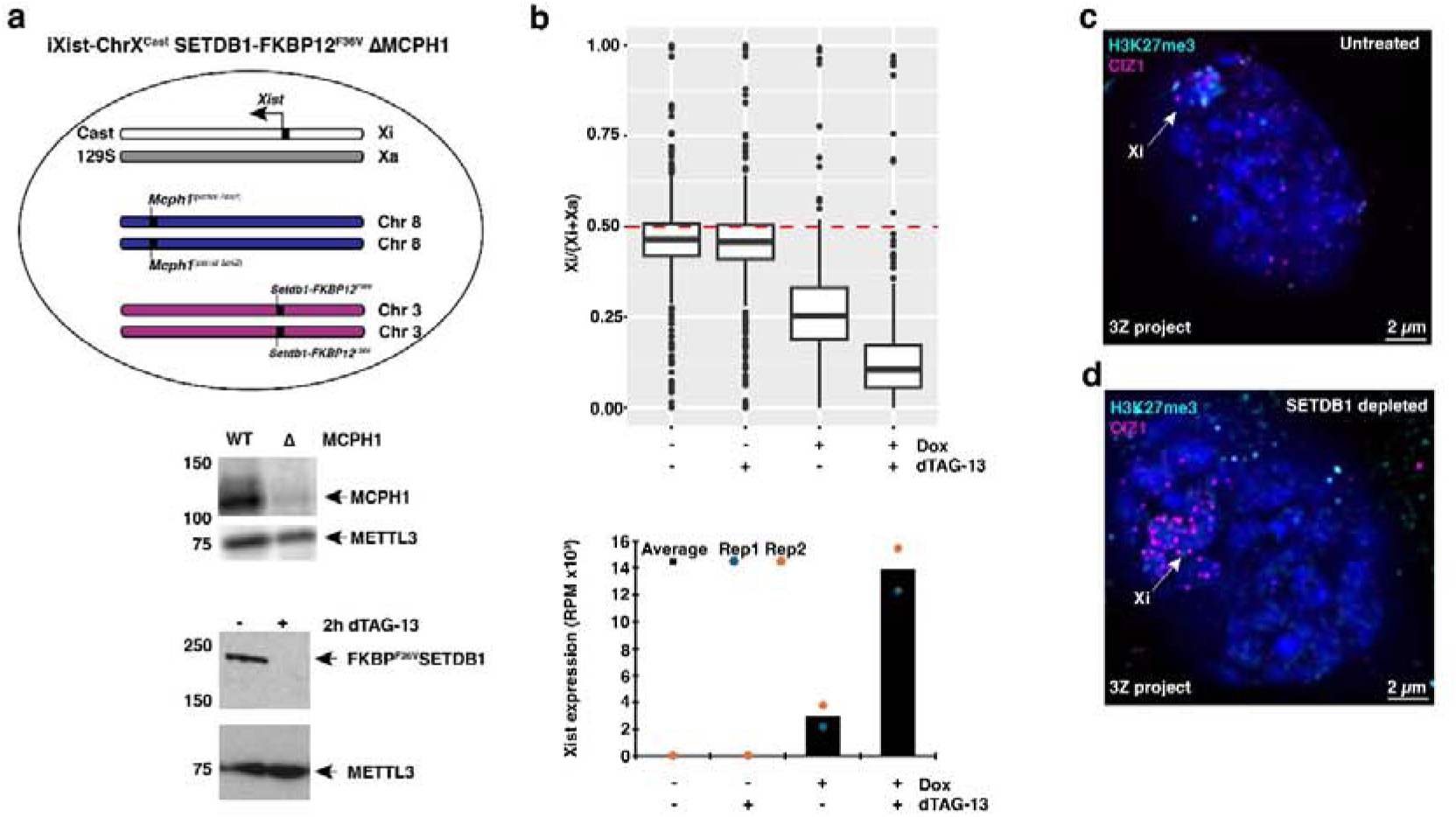
Analysis of XCI in iXist-ChrX^Cast^ ΔMCPH1 mESCs after SETDB1 depletion. **a.** Schematic illustrating interspecies *Mus domesticus (129S) x Mus castaneus (Cast)* iXist-ChrX^Cast^ SETDB1-FKBP12^F36V^ XX mouse ESC line into which a homozygous deletion was engineered into *Mcph1* on chromosome 8. Xist expression was induced from the *Cast* X chromosome via rtTA in response to doxycycline (dox) as previously described^15,20^. Western blots show WT and ΔMCPH1 nuclear extracts probed with anti-MCPH1 with anti-METTL3 loading control (top), and whole cell extracts from WT and SETDB1-FKBP12^F36V^ ESCs +/- 2 h dTAG-13 treatment, probed with anti-SETDB1 with anti-METTL3 loading control (bottom). **b.** ChrRNA-seq analysis showing silencing of Xi genes is accelerated when Xist expression is induced in the absence of SETBD1 in iXist-ChrX^Cast^ SETBD1-FKBP12^F36V^ ΔMCPH1 mESCs as is observed in the presence of MCPH1 (top panel)^15^. Corresponding Xist expression levels determined from ChrRNA-seq in iXist-ChrX^Cast^ SETBD1-FKBP12^F36V^ ΔMCPH1 mESCs (average of two replicas) in the presence and absence of doxycycline and/or removal of SETDB1 by dTAG-13 treatment (bottom panel). n=464 genes. IF for H3K27me3 and CIZ1 in **c.** untreated iXist-ChrX^Cast^ SETBD1-FKBP12^F36V^ ΔMCPH1 cell with condensed chromosomes and **d**. after treatment with dTAG-13 to deplete SETDB1. Images are deconvolved widefield 3z maximum intensity projections. DNA was counterstained with DAPI (blue).

**Extended Data Figure 6.**
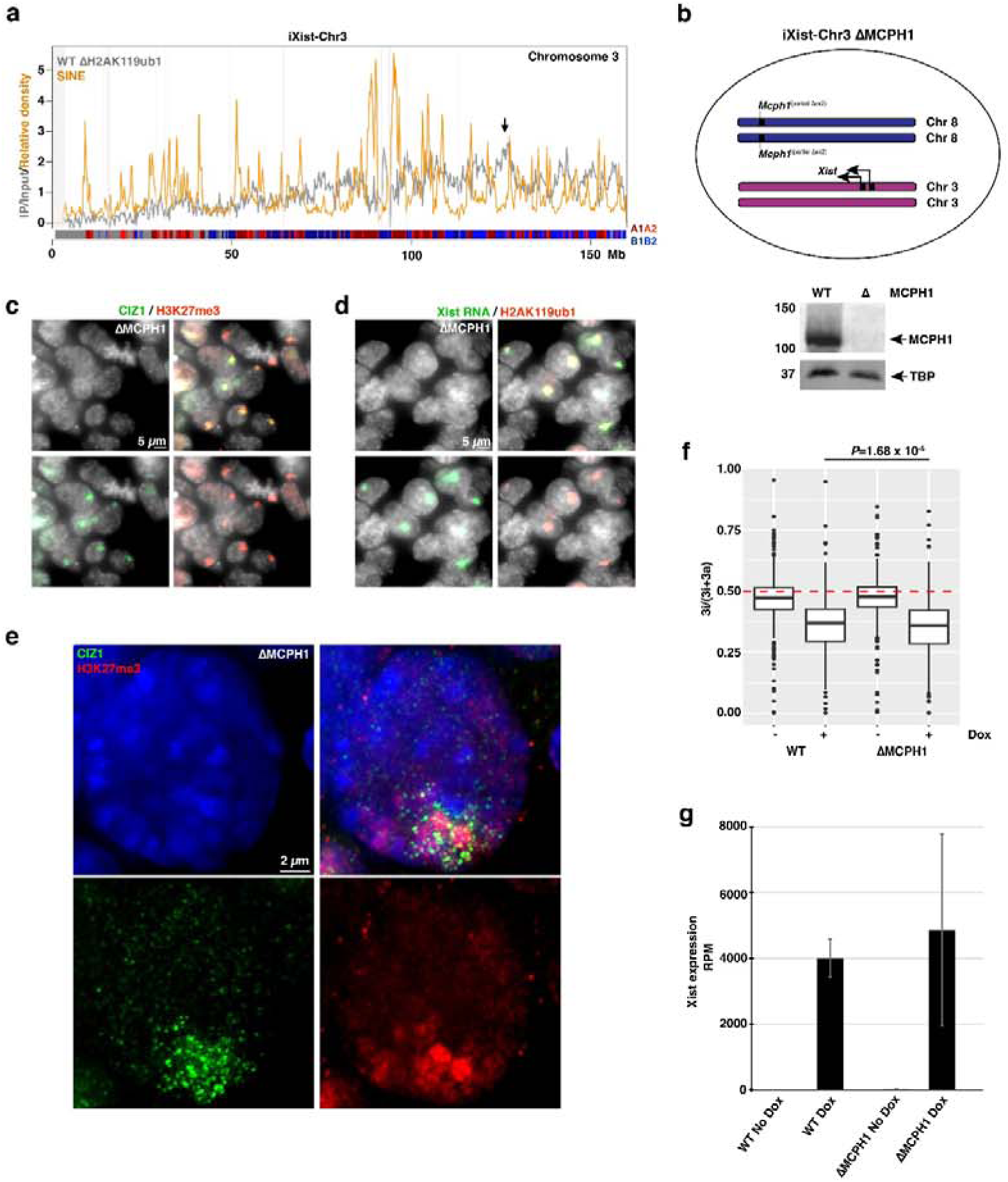
Analysis of XCI in ΔMCPH1 iXist-Chr3 cell line. **a.** Distribution of H2AK119ub1 gain relative to SINE density after Xist induction in iXist-Chr3 XY mESCs. Arrow indicates the chromosome 3 Xist transgene integration site. **b**. Schematic illustrating the transgenic iXist-Chr3 XY mouse ESC line into which a homozygous deletion was engineered into *Mcph1* on chromosome 8. Transgenic multicopy Xist expression was induced in response to doxycycline (dox) as previously described^20^ (upper panel). Western blot of nuclear extracts from WT and MCPH1 null cells, probed with anti-MCPH1 and anti-TBP as a loading control (lower panel). **c.** Widefield image of H3K27me3 and CIZ1 IF in ΔMCPH1 iXist-Chr3 mESCs. **d.** Widefield image of Xist RNA and CIZ1 immuno-FISH in ΔMCPH1 iXist-Chr3 mESCs. **e.** H3K27me3 and CIZ1 IF in ΔMCPH1 iXist-Chr3 mESCs; maximum intensity projection of deconvolved widefield image; DNA was counterstained with DAPI (blue). **f.** ChrRNA-seq showing chromosome 3 gene silencing is significantly accelerated (paired T-test) when Xist expression is induced in ΔMCPH1 iXist-Chr3 compared to iXist-Chr3 mESCs^20^. n=580 genes. **g.** Xist expression levels determined from ChrRNA-seq in iXist-Chr3 mESCs in the presence^20^ and absence of MCPH1 and +/-doxycycline induction (average shown of four replica experiments).

**Extended Data Figure 7.**
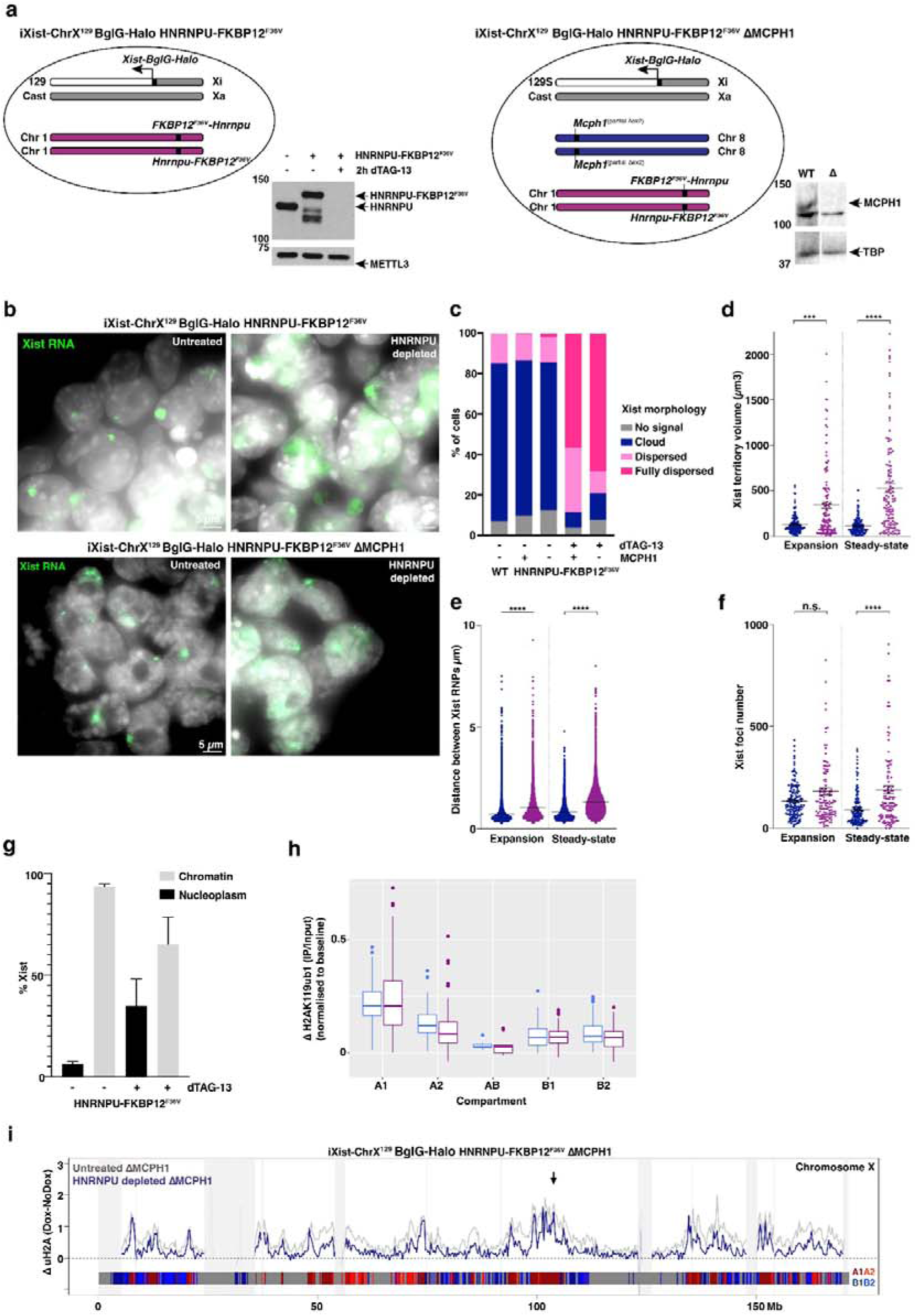
Analysis of XCI in ΔMCPH1 after HNRNPU depletion. **a.** Schematics illustrating (left) interspecific *Mus domesticus (129S) x Mus castaneus (Cast)* iXist-ChrX^129^ BglG-Halo ESC line in which FKBP12^F36V^ was engineered into the endogenous *Hnrnpu* locus on chromosome 1. Xist expression was induced from the *129S* X chromosome as described^20^. Western blot of nuclear extracts from WT and HNRNPU-FKBP12^F36V^ ESCs +/- 2 h dTAG-13 treatment, probed with anti-HNRNPU and anti-METTL3 loading control. Subsequent engineering of iXist-ChrX^129^ BglG-Halo HNRNPU-FKBP12^F36V^ mESCs with a homozygous deletion of *Mcph1* on chromosome 8 (right). Western blot of nuclear extracts from WT and ΔMCPH1 probed with anti-MCPH1 and anti-TBP loading control. **b**. Xist RNA FISH (green) and DNA stained with DAPI (grey), showing that HNRNPU-deficient XX mESCs exhibit a dispersed Xist cloud (maximum intensity projection). **b.** Quantification of Xist RNA FISH cloud morphology in WT, and HNRNPU-FKBP12^F36V^ cells +/- 26 h dTAG-13 treatment and 24 h dox induction of Xist, n= 200. **d.** RNA-SPLIT analysis of Xist territory volume *** p = 0.0001, **** p <0.0001, Blue = WT, Purple = HNRNPU-depleted (n = 161 cells WT expansion, 133 WT steady-state, 122 HNRNPU-FKBP12^F36V^ expansion, 116 HNRNPU-FKBP12^F36V^ steady-state. Expansion phase refers to analysis after 1.5 h doxycycline induction and Steady-state phase to analysis after 24 h doxycycline induction. **e.** RNA-SPLIT analysis of distance between Xist foci (nm), **** p <0.0001, Blue = WT, Purple = HNRNPU-depleted (n = 18,042 foci WT expansion, 4,988 WT steady-state, n = 13,589 HNRNPU-FKBP12^F36V^ expansion, n= 16,469 HNRNPU-FKBP12^F36V^ steady-state). **f**. RNA-SPLIT analysis of Xist foci number, n.s = not significant, **** p <0.0001, Blue =WT. Purple =HNRNPU-depleted. (n = 153 cells WT expansion, 133 WT steady-state, 121 HNRNPU-FKBP12^F36V^ expansion, 117 HNRNPU-FKBP12^F36V^ steady-state. **g.** qRT-PCR analysis of Xist RNA distribution in soluble nucleoplasmic and insoluble chromatin fractions +/- 22 h dTAG-13 treatment and 20 h Xist induction, normalised to actin (3 replicates, 2 biological replicates). **h**. Box plot analysis showing Xi gain of H2AK119ub1 within different chromatin sub-compartments in untreated HNRNPU-FKBP12^F36V^ mESCs (blue) vs HNRNPU-depleted cells (purple; data from two replicas of each condition as above, two high-value data points not included on the chart for A1 HNRNPU-depleted cells for conciseness). Data baseline-normalised rather than absolute H2AK119ub1 levels to better distinguish changes in compartment distribution. **i**. Xi gain of H2AK119ub1 determined from nChIP in iXist-ChrX^129^ BglG-Halo HNRNPU-FKBP12^F36V^ ΔMCPH1 mESCs (blue line, two merged replicas). H2AK119ub1 gain is reduced across the X chromosome when HNRNPU is depleted (blue line, two merged replicas). Blocked out panels (light grey) represent regions/genomic bins where the input signal is significantly different (three standard deviations) from its mean value as computed across the X chromosome (see Materials and Methods). Chromosome sub-compartments depicted in lower bar: A1 dark red, A2 light red, B1 dark blue, B2 light blue, dark grey uncertain compartment assignment. The arrow indicates the location of the Xist gene.

## Supplementary Data

### Supplementary Movie 1

3D rendering zooming in on Xi from 3D-SIM image z-stack for Xist RNA FISH (green) of G2 iXist-ChrX^129^ ΔMCPH1 cell. DNA is counterstained with DAPI (blue).

### Supplementary Movie 2

3D rendering zooming in on Xi from 3D-SIM image z-stack for Xist RNA FISH (green) of G2 C127 ΔMCPH1 cell. DNA is counterstained with DAPI (blue).

### Supplementary Movie 3

3D rendering zooming in on Xi from 3D-SIM image z-stack for Xist RNA FISH (green) of WT C127 prophase cell. DNA is counterstained with DAPI (blue).

## References

1. Lyon, M. F. Gene Action in the X-chromosome of the Mouse (Mus musculus L.). Nature 190, 372–373 (1961).

2. Brockdorff, N. et al. The product of the mouse Xist gene is a 15 kb inactive X-specific transcript containing no conserved ORF and located in the nucleus. Cell 71, 515–526 (1992).

3. Brown, C. J. et al. A gene from the region of the human X inactivation centre is expressed exclusively from the inactive X chromosome. Nature 349, 38–44 (1991).

4. Brown, C. J. et al. The human XIST gene: Analysis of a 17 kb inactive X-specific RNA that contains conserved repeats and is highly localized within the nucleus. Cell 71, 527–542 (1992).

5. Penny, G. D., Kay, G. F., Sheardown, S. A., Rastan, S. & Brockdorff, N. Requirement for Xist in X chromosome inactivation. Nature 379, 131–137 (1996).

6. Lee, J. T. & Jaenisch, R. Long-range cis effects of ectopic X-inactivation centres on a mouse autosome. Nature 386, 275–279 (1997).

7. Loda, A., Collombet, S. & Heard, E. Gene regulation in time and space during X-chromosome inactivation. Nat. Rev. Mol. Cell Biol. 23, 231–249 (2022).

8. Brockdorff, N., Bowness, J. S. & Wei, G. Progress toward understanding chromosome silencing by Xist RNA. Genes Dev. 34, 733–744 (2020).

9. Keniry, A. & Blewitt, M. E. Chromatin-mediated silencing on the inactive X chromosome. Development 150, dev201742 (2023).

10. Bowness, J. S. et al. Xist-mediated silencing requires additive functions of SPEN and Polycomb together with differentiation-dependent recruitment of SmcHD1. Cell Rep. 39, 110830 (2022).

11. Hasegawa, Y. et al. The Matrix Protein hnRNP U Is Required for Chromosomal Localization of Xist RNA. Dev. Cell 19, 469–476 (2010).

12. Ridings-Figueroa, R. et al. The nuclear matrix protein CIZ1 facilitates localization of Xist RNA to the inactive X-chromosome territory. Genes Dev. 31, 876–888 (2017).

13. Engreitz, J. M. et al. The Xist lncRNA exploits three-dimensional genome architecture to spread across the X chromosome. Science 341, 1237973 (2013).

14. Wei, G. et al. m6A and the NEXT complex direct Xist RNA turnover and X-inactivation dynamics. Nat. Struct. Mol. Biol. 32, 2242–2251 (2025).

15. Almeida, M., Wei, G., Cawte, A., Nesterova, T. & Brockdorff, N. SETDB1 and HUSH modulate Xist RNA levels during establishment of X chromosome inactivation. Nat. Commun. in press,.

16. Ding, M. et al. A biophysical basis for the spreading behavior and limited diffusion of Xist. Cell 188, 978–997.e25 (2025).

17. Pandya-Jones, A. et al. A protein assembly mediates Xist localization and gene silencing. Nature 587, 145–151 (2020).

18. Houlard, M. et al. MCPH1 inhibits Condensin II during interphase by regulating its SMC2-Kleisin interface. eLife 10, e73348 (2021).

19. Rodermund, L. et al. Time-resolved structured illumination microscopy reveals key principles of Xist RNA spreading. Science 372, eabe7500 (2021).

20. Nesterova, T. B. et al. Systematic allelic analysis defines the interplay of key pathways in X chromosome inactivation. Nat. Commun. 10, 3129 (2019).

21. Smeets, D. et al. Three-dimensional super-resolution microscopy of the inactive X chromosome territory reveals a collapse of its active nuclear compartment harboring distinct Xist RNA foci. Epigenetics Chromatin 7, 8 (2014).

22. Markaki, Y. et al. Xist nucleates local protein gradients to propagate silencing across the X chromosome. Cell 184, 6174–6192.e32 (2021).

23. Rao, S. S. P. et al. A 3D Map of the Human Genome at Kilobase Resolution Reveals Principles of Chromatin Looping. Cell 159, 1665–1680 (2014).

24. Liu, J., Ali, M. & Zhou, Q. Establishment and evolution of heterochromatin. Ann. N. Y. Acad. Sci. 1476, 59–77 (2020).

25. Brown, J. M., De Ornellas, S., Parisi, E., Schermelleh, L. & Buckle, V. J. RASER-FISH: non-denaturing fluorescence in situ hybridization for preservation of three-dimensional interphase chromatin structure. Nat. Protoc. 17, 1306–1331 (2022).

26. Lieberman-Aiden, E. et al. Comprehensive Mapping of Long-Range Interactions Reveals Folding Principles of the Human Genome. Science 326, 289–293 (2009).

27. Gibcus, J. H. et al. A pathway for mitotic chromosome formation. Science 359, eaao6135 (2018).

28. Jachowicz, J. W. et al. Xist spatially amplifies SHARP/SPEN recruitment to balance chromosome-wide silencing and specificity to the X chromosome. Nat. Struct. Mol. Biol. 29, 239–249 (2022).

29. Dror, I. et al. XIST directly regulates X-linked and autosomal genes in naive human pluripotent cells. Cell 187, 110–129.e31 (2024).

30. Yao, S., Jeon, Y., Kesner, B. & Lee, J. T. Xist RNA binds select autosomal genes and depends on Repeat B to regulate their expression. eLife 13, RP101197 (2025).

31. Nabet, B. et al. The dTAG system for immediate and target-specific protein degradation. Nat. Chem. Biol. 14, 431–441 (2018).

32. Chu, C. et al. Systematic Discovery of Xist RNA Binding Proteins. Cell 161, 404–416 (2015).

33. McHugh, C. A. et al. The Xist lncRNA interacts directly with SHARP to silence transcription through HDAC3. Nature 521, 232–236 (2015).

34. Sharp, J. A. et al. Role of the SAF-A/HNRNPU SAP domain in X chromosome inactivation, nuclear dynamics, transcription, splicing, and cell proliferation. PLoS Genet. 21, e1011719 (2025).

35. Duthie, S. M. et al. Xist RNA exhibits a banded localization on the inactive X chromosome and is excluded from autosomal material in cis. Hum. Mol. Genet. 8, 195–204 (1999).

36. Walter, J., Schermelleh, L., Cremer, M., Tashiro, S. & Cremer, T. Chromosome order in HeLa cells changes during mitosis and early G1, but is stably maintained during subsequent interphase stages. J. Cell Biol. 160, 685–697 (2003).

37. Moindrot, B. et al. A Pooled shRNA Screen Identifies Rbm15, Spen, and Wtap as Factors Required for Xist RNA-Mediated Silencing. Cell Rep. 12, 562–572 (2015).

38. Demmerle, J. et al. Strategic and practical guidelines for successful structured illumination microscopy. Nat. Protoc. 12, 988–1010 (2017).

39. Ball, G. et al. SIMcheck: a Toolbox for Successful Super-resolution Structured Illumination Microscopy. Sci. Rep. 5, 15915 (2015).

40. Matsuda, A., Schermelleh, L., Hirano, Y., Haraguchi, T. & Hiraoka, Y. Accurate and fiducial-marker-free correction for three-dimensional chromatic shift in biological fluorescence microscopy. Sci. Rep. 8, 7583 (2018).

41. Kraus, F. et al. Quantitative 3D structured illumination microscopy of nuclear structures. Nat. Protoc. 12, 1011–1028 (2017).

42. Langmead, B. & Salzberg, S. L. Fast gapped-read alignment with Bowtie 2. Nat. Methods 9, 357–359 (2012).

43. Dobin, A. et al. STAR: ultrafast universal RNA-seq aligner. Bioinformatics 29, 15–21 (2013).

44. Liao, Y., Smyth, G. K. & Shi, W. featureCounts: an efficient general purpose program for assigning sequence reads to genomic features. Bioinformatics 30, 923–930 (2014).

45. Li, H. et al. The Sequence Alignment/Map format and SAMtools. Bioinformatics 25, 2078–2079 (2009).

46. Quinlan, A. R. & Hall, I. M. BEDTools: a flexible suite of utilities for comparing genomic features. Bioinforma. Oxf. Engl. 26, 841–842 (2010).

47. Robinson, J. T. et al. Integrative genomics viewer. Nat. Biotechnol. 29, 24–26 (2011).

48. Bowness, J. S. et al. Xist-mediated silencing requires additive functions of SPEN and Polycomb together with differentiation-dependent recruitment of SmcHD1. Cell Rep. 39, 110830 (2022).

49. Lawrence, M. et al. Software for Computing and Annotating Genomic Ranges. PLoS Comput. Biol. 9, e1003118 (2013).

50. Gel, B. & Serra, E. karyoploteR: an R/Bioconductor package to plot customizable genomes displaying arbitrary data. Bioinformatics 33, 3088–3090 (2017).

51. Mudge, J. M. et al. GENCODE 2025: reference gene annotation for human and mouse. Nucleic Acids Res. 53, D966–D975 (2025).

52. Bauer, M. et al. Chromosome compartments on the inactive X guide TAD formation independently of transcription during X-reactivation. Nat. Commun. 12, 3499 (2021).

53. Wang, C.-Y., Jégu, T., Chu, H.-P., Oh, H. J. & Lee, J. T. SMCHD1 Merges Chromosome Compartments and Assists Formation of Super-Structures on the Inactive X. Cell 174, 406–421.e25 (2018).

54. Brown, A. C. et al. Comprehensive epigenomic profiling reveals the extent of disease-specific chromatin states and informs target discovery in ankylosing spondylitis. Cell Genomics 3, 100306 (2023).

55. Magoč, T. & Salzberg, S. L. FLASH: fast length adjustment of short reads to improve genome assemblies. Bioinformatics 27, 2957–2963 (2011).

56. Wingett, S., et al. HiCUP: pipeline for mapping and processing Hi-C data. F1000Research 4, 1310 (2015).

57. Telenius, J., The WIGWAM Consortium & Hughes, J. R. NGseqBasic - a single-command UNIX tool for ATAC-seq, DNaseI-seq, Cut-and-Run, and ChIP-seq data mapping, high-resolution visualisation, and quality control. Preprint at 10.1101/393413 (2018).

58. Krueger, F. & Andrews, S. R. SNPsplit: Allele-specific splitting of alignments between genomes with known SNP genotypes. F1000Research 5, 1479 (2016).

59. Imakaev, M. et al. Iterative correction of Hi-C data reveals hallmarks of chromosome organization. Nat. Methods 9, 999–1003 (2012).

60. Open2C et al. Cooltools: Enabling high-resolution Hi-C analysis in Python. PLOS Comput. Biol. 20, e1012067 (2024).

61. Servant, N., et al. HiC-Pro: an optimized and flexible pipeline for Hi-C data processing. Genome Biol. 16, 259 (2015).

62. Jin, Y., Tam, O. H., Paniagua, E. & Hammell, M. TEtranscripts: a package for including transposable elements in differential expression analysis of RNA-seq datasets. Bioinformatics 31, 3593–3599 (2015).

63. Lu, J. Y. et al. Homotypic clustering of L1 and B1/Alu repeats compartmentalizes the 3D genome. Cell Res. 31, 613–630 (2021).

64. Lafontaine, D. L., Yang, L., Dekker, J. & Gibcus, J. H. Hi-C 3.0: Improved Protocol for Genome-Wide Chromosome Conformation Capture. Curr. Protoc. 1, e198 (2021).

65. Durand, N. C. et al. Juicer Provides a One-Click System for Analyzing Loop-Resolution Hi-C Experiments. Cell Syst. 3, 95–98 (2016).

66. Ramírez, F. et al. High-resolution TADs reveal DNA sequences underlying genome organization in flies. Nat. Commun. 9, 189 (2018).

